# Physical Activity Induces Nucleus Accumbens Genes Expression Changes Preventing Chronic Pain Susceptibility Promoted by High-Fat Diet and Sedentary Behavior in Mice

**DOI:** 10.1101/768309

**Authors:** Arthur de Freitas Brandão, Ivan José Magayewski Bonet, Marco Pagliusi, Gabriel Gerardini Zanetti, Nam Pho, Cláudia Herrera Tambeli, Carlos Amilcar Parada, André Schwambach Vieira, Cesar Renato Sartori

## Abstract

High-fat diet (HFD)-induced obesity was reported to increase pain behavior independent of obesity status in rats, whereas weight loss interventions such as voluntary physical activity (PA) for adults with overweight or obesity was reported to promote pain reduction in humans with chronic pain (CP). However, is unknown whether an HFD and sedentary (SED) behavior is underlying to CP susceptibility and whether voluntary PA can prevent it. Moreover, differential gene expression in the nucleus accumbens (NAc) is considered to play a crucial role in CP susceptibility. The present study used an adapted model of the inflammatory prostaglandin E2 (PGE)-induced persistent hyperalgesia (PH-ST) protocol for mice, an HFD, and a voluntary PA paradigm to test these hypotheses. In addition, we performed a transcriptome in the NAc and a gene ontology enrichment tools to investigate the differential gene expression and identify the biological processes associated with CP susceptibility tested here. Our results demonstrated that HFD and sedentary behavior promoted CP susceptibility, which in turn was prevented by voluntary PA, even when the animals were fed an HFD. Transcriptome in the NAc found 2,204 differential expression genes related CP susceptibility promoted by HFD and sedentary behavior and prevented by voluntary PA. The gene ontology enrichment revealed 41 biological processes implicated in CP susceptibility. Analyzing collectively those biological processes, our results suggested that genes related to metabolic and mitochondria stress were up-regulated in the CP susceptibility group, whereas genes related to neuroplasticity and axonogenesis were up-regulated in the CP prevented group. These findings provide pieces of evidence that an HFD and sedentary behavior promoted gene expression changes in the NAc related to neurodegeneration and those changes were also underlying to CP susceptibility. Additionally, our findings confirmed other findings supporting the crucial role of voluntary PA to prevent CP susceptibility and add novel insights of differential gene expression in the NAc related to neuroplasticity.

## 1. Introduction

Obesity and chronic pain (CP) are two highly prevalent conditions associated with a modern lifestyle in both developed and developing countries. A meta-analysis study from 1975 to 2016 showed an increase in overweight and obesity prevalence worldwide and the authors argued that an unhealthy nutritional transition (*i.e.* childhood to adulthood) and an increase of nutrient-poor with high energy-dense foods (*e.g.* fat and sugar) lead to weight gain (Abarca-Gómez et al., 2017). At the same time, epidemiological studies indicate an increase of humans with CP in recent years (Torrance et al., 2006; Van Hecke et al., 2013, 2014) with an economic cost between $560 to 635 billion annually in the United States alone (Gaskin and Richard, 2012) and with major impacts on individual and social life (Breivik et al., 2013; Mayer et al., 2019).

A cross-sectional study from low- and middle-income countries showed that sedentary behavior is strongly related to obesity and CP and suggested that interventions focusing on reducing the sedentary behavior should be considered interventions for those chronic conditions (Koyanagi et al., 2018). Furthermore, weight loss interventions such as voluntary physical activity (PA) for adults with overweight or obesity promote significant pain reduction in humans with CP (Cooper et al., 2018a). In fact, the amount of evidence related to the benefits of PA as a treatment for individuals with CP are widely reported in the literature (Geneen et al., 2017; Lima et al., 2017), as well as for individuals with obesity (Paley and Johnson, 2016). However, these studies did not investigate the interaction between high-fat diet (HFD)-induce obesity, sedentary behavior, voluntary PA, or weight lost in CP susceptibility, neither a potential alteration in the central nervous system underlying those variables. Several studies suggested brain neuroplasticity is underlying to comorbidity of CP (Apkarian et al., 2009; Doan et al., 2015; Mansour et al., 2014) and the nucleus accumbens (NAc) has been investigated as having a critical role in modulating CP in humans and animal models (Benarroch, 2016; Martikainen et al., 2015; Ren et al., 2015; Schwartz et al., 2017).

Transcriptomic studies to describe differential gene expression and biological processes related to CP susceptibility have been used to reveal novel insights (Starobova et al., 2018). For instance, to investigate migraine-associated hyperalgesia, a recent transcriptome study of mice NAc and trigeminal ganglia showed gene network dysregulation and biological processes alteration in both areas (Jeong et al., 2018). Through a gene ontology enrichment tools, the authors identified crucial up-regulated biological processes in both areas related to hyperalgesia, such as amino acid transmembrane, anion transmembrane transport, and amino acid transport (Jeong et al., 2018). Other CP transcriptome study of mice NAc, medial prefrontal cortex (mPFC), and periaqueductal gray (PAG) showed that gene expression changes in these brain networks were strongly related with other comorbidities, including neuropathic pain, depression, and stress (Descalzi et al., 2017).

Additionally, Jayaraman et al. (2014) demonstrate glial cultures generated from high-fat-fed animals exhibit reduced survival, poorer neurite outgrowth, and evidence of nerve damage in peripheral nervous system. Although these alterations have been identified from primary neurons co-cultured with glial cultures, investigation of biological processes related to metabolic and mitochondrial stress in central nervous system (*i.e.* NAc) associated with an HFD-induced obesity, CP susceptibility, and voluntary PA remain elusive. Thus, a general understanding the complex landscape of biological processes underlying CP susceptibility induced by an HFD and sedentary behavior, as well as the role of voluntary PA in preventing CP susceptibility through a transcriptome of mice NAc can shed some lights and reveal novel interesting insights for science.

The first objective of this study was investigating whether HFD and sedentary behavior promote CP susceptibility and whether voluntary PA can prevent it. Our hypothesis was that an HFD and sedentary behavior would lead to CP susceptibility and voluntary PA would prevent it. Our second objective was describing the differential gene expression in the NAc related to CP susceptibility promoted by an HFD and sedentary behavior and prevented by voluntary PA. Finally. we also applied gene ontology enrichment tools to describe the biological processes related to CP susceptibility promoted by an HFD and sedentary behavior and prevented by voluntary PA.

## 2. Methods

### 2.1 Animal

To test our hypothesis, eighty male C57BL/6JUnib mice at the age of 4 weeks were used from the Multidisciplinary Center for Biological Investigation on Laboratory Animal Science of University of Campinas (CEMIB - UNICAMP). The mice were housed in a temperature-controlled room (21±1°C) on a 12 hour light/dark cycle (lights on at 7 AM) with *ad libitum* access to food and water. Before the beginning of the experiment, all mice were acclimated to the vivarium in collective cages (5 mice per cage) for two weeks. At six weeks old, the mice were randomly moved to an individual cage through the end of the experiment (at 18 weeks old). All procedures were reviewed and approved in accordance to animal experimentation of Brazilian federal law (n° 11.794 October 08, 2008) and by the animal ethics committee at the University of Campinas (CEUA-UNICAMP) under protocols number 4243-1. All efforts were made to reduce the number of animals used and minimize animal suffering.

### 2.2 Experimental Design

At six weeks old, the mice were randomly separated in two groups: one was fed a standard diet (SD; *n* = 40) and the other a high-fat diet (HFD; *n* = 40) for 12 weeks (**Figure 1**). At 12 weeks old, it means after six weeks feeding one of the experimental diets, the SD and HFD groups were again randomly subdivided into four groups: i) sedentary/standard diet (SED-SD; *n* = 20) and, ii) sedentary/high-fat diet (SED-HFD; *n* = 20) groups, which were placed in standard cages without a running wheel (RW) access; iii) physically active/standard diet (PA-SD; *n* = 20) and iv) physically active/high-fat diet (PA-HFD; *n* = 20) groups, which were placed in cages with free access to RW for six weeks. At 16 weeks old, each group was subdivided into two sub-groups, one of those sub-groups was submitted to PGE2-induced persistent hyperalgesia with a short-term (PH-ST) induction protocol and the other group was submitted to a saline administration as a PGE2 control group (details in 2.5 sessions). Thus, at the end of the experimental design, we had eight experimental groups (*n* = 10, each group) according to **Table 1**.

**Figure 1:**
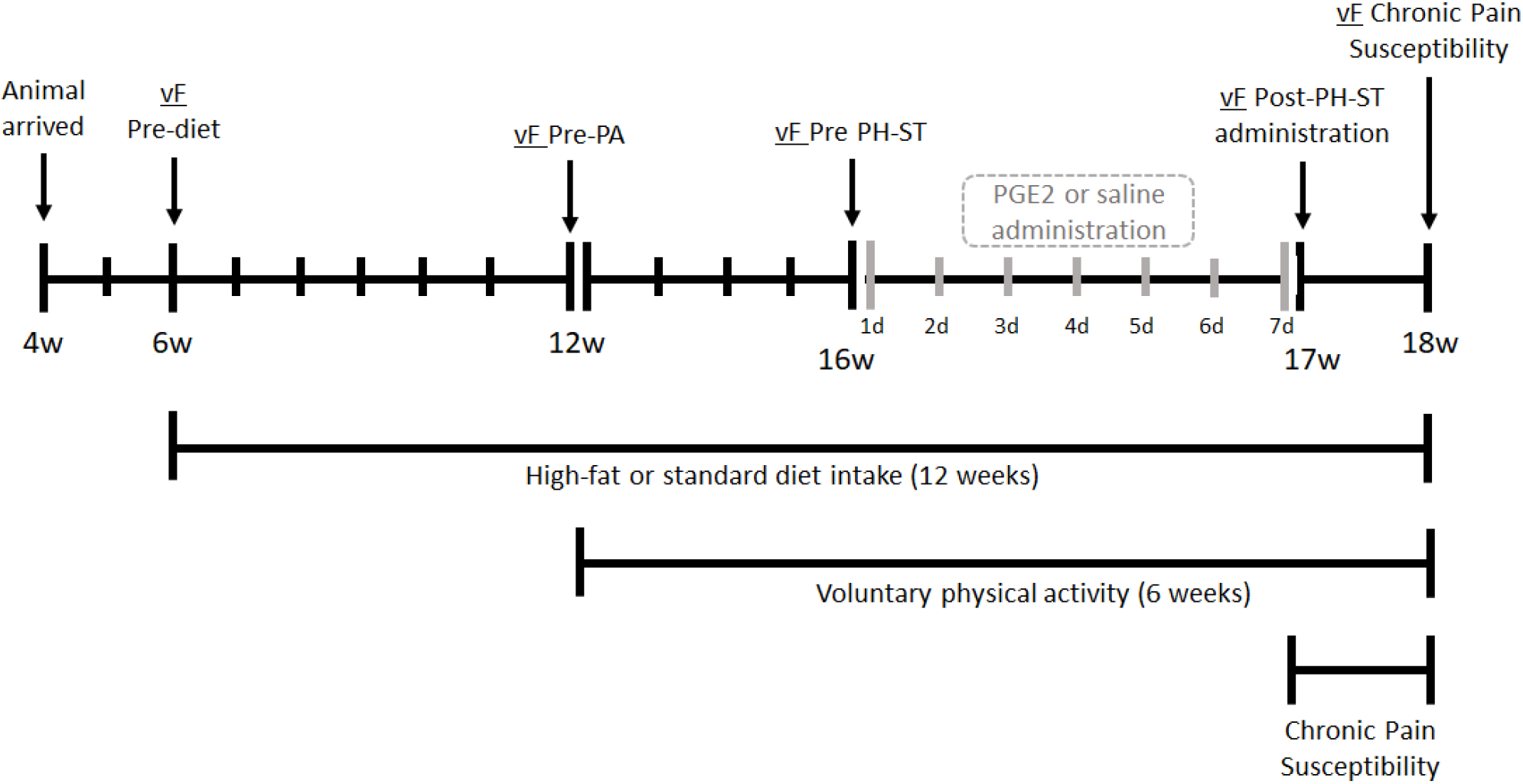
Experimental design. At the age of the 4 weeks, the mice arrived and for two weeks they acclimated with a new vivarium. At 6 weeks old, the mice were subdivided into two groups and each group was exposed to one of the different diets (standard or high fat) for 12 weeks and each group was submitted to the first mechanical nociceptive threshold through vF Pre-diet test. At 12 weeks old, the mice were subdivided in four groups, being that, two groups (PA-SD and PA-HFD) had a free running wheel accesses for voluntary physical activity (PA) and another two groups (SED-SD and SED-HFD) remain sedentary (SED) in a standard cage for six weeks, and all four groups were submitted to vF Pre-PA test. At 16 weeks old, each group was submitted to vF Pre-PH-ST test before starting the administration of PGE2 (PH-ST protocol) or saline. At 17 weeks old, all eight groups were submitted to vF Post-PH-ST administration test. At 18 weeks old, all eight groups were submitted to vF chronic pain (CP) susceptibility test and were euthanized. vF = von Frey, PA = physical activity, PH-ST = persistent hyperalgesia short-term protocol, PGE2 = prostaglandin E2, w = weeks old age.

**Table 1:** Groups identification and description, and sample number for all stages of experiment design. SD = standard diet, HFD = high-fat diet, S = sedentary, PA = physically active, PGE = prostaglandin E2 and SAL = saline. n1 = number of mice per group initially used; n2 = number of mice per group for all behavior’s analysis; n3 = number of mice per group for transcriptome analysis.

| Groups | Description | *n1, n2, n3* |
| --- | --- | --- |
| SED-SD-SAL | Sedentary mice fed a standard diet and received saline | 10-8-5 |
| SED-SD-PGE | Sedentary mice fed a standard diet and received PGE | 10-9-5 |
| PA-SD-SAL | Physically active mice fed a standard diet and received saline | 10-8-5 |
| PA-SD-PGE | Physically active mice fed a standard diet and received PGE | 10-9-5 |
| SED-HFD-SAL | Sedentary mice fed a high-fat diet and received saline | 10-8-5 |
| SED-HFD-PGE | Sedentary mice fed a high-fat diet and received PGE | 10-9-5 |
| PA-HFD-SAL | Physically active mice fed a high-fat diet and received saline | 10-9-5 |
| PA-HFD-PGE | Physical active mice fed a high-fat diet and received PGE | 10-9-5 |

Furthermore, we performed five measures of mechanical nociceptive threshold through an electronic von-Frey (vF) test. The first vF test was performed before each group was exposed to different diets (vF Pre-diet) at 6 weeks old. The second vF test was performed before each group had free access to RW for voluntary PA (vF Pre-PA) at 12 weeks old. The third vF test was performed 3 hours after the first dose of the PGE2-induced the PH-ST protocol or saline administration (vF Post-PH-ST first dose) at 16 weeks and one day old. The fourth vF test was performed one day after the end of PGE2-induced PH-ST protocol or saline administration (vF Post-PGE2 administration) at 17 weeks old. The fifth vF test was performed seven days after the end of the PH-ST protocol, at 18 weeks old (**Figure 1**). During CP susceptibility period, from 17 until 18 weeks old, the mice were not manipulated.

### 2.3 Body Mass, Caloric Intake and Diets

Body mass and food intake were recorded weekly, always at the same weekday and at the same time of day (1 PM ± 1 hour, light cycle). The SD (3.080 kcal/g) was composed by 11.7% kcal/g from lipid, 28.5% kcal/g from protein and, 59.7% kcal/g from carbohydrate (Nuvilab-Quimtia S/A. Paraná, Brazil) **Table 2**. The HFD (5.439 kcal/g) was composed by 58.2% kcal/g from lipid (51.61% from lard and 6.61% kcal/g from soybean oil), 14.9% kcal/g from protein and, 26.8% kcal/g from carbohydrate provided by Laboratory of Nutritional Genomic from the School of Applied Science of the University of Campinas (Cintra et al., 2012).

**Table 2:** Distribution of the macronutrients and caloric equivalent (kcal/g) of diets.

| Macronutrients | Standard diet<br>(kcal%) | High-fat diet<br>(kcal%) |
| --- | --- | --- |
| Lipid | 11.7 | 58.2 |
| Protein | 28.5 | 14.9 |
| Carbohydrate | 59.7 | 26.8 |
| **TOTAL** | **3.080 kcal/g** | **5.439 kcal/g** |

### 2.4 Adipose Tissue Dissection

To investigate the amount and the effect of voluntary PA on adipose tissue, we dissected and immediately weigh the epididymal and retroperitoneal adipose tissue as described by de Jong et al. (2015). Afterward, we snap-froze both adipose tissues in liquid nitrogen and stored them in a −80° C freezer. The data of each adipose tissue (in grams) was normalized with total body mass (in grams) and was presented by the mean percentage and standard error.

### 2.5 Prostaglandin E2-Induced Persistent Hyperalgesia Short-Term Protocol

To test CP susceptibility, we employed an adapted model of the inflammatory prostaglandin E2 (PGE2)-induced persistent hyperalgesia (PH-ST) protocol previously described by Ferreira et al. (1990) and Villarreal et al. (2009). Briefly, the original model showed that 14 successive days of intraplantar injection of PGE2 was sufficient to induce persistent hyperalgesia in rodents for more than thirty days after the last PGE2 injection (Ferreira et al., 1990; Villarreal et al., 2009). However, here we used a PH-ST protocol with only seven successive days of intraplantar injection of PGE2, which was not enough to produce persistent hyperalgesia (Dias et al., 2015; Ferreira et al., 1990). This model was advantageous compare to others’ CP models because further pharmacological studies revealed a key role of non-inflammatory agents in the chronification process, such as PKA (protein kinase A), PKCε (protein kinase C isoform ε), AC (adenylyl cyclase) (Sachs et al., 2009; Villarreal et al., 2009) and NF-κB (nuclear factor kappa-B) (Souza et al., 2015). PGE2 and saline (SAL) administration was made through a hypodermic 26-gauge needle coupled at a 50μL Hamilton syringe to administer 18μL [90ng] of PGE2 (Sigma – Aldrich®, St. Louis, MO, USA) or 18μL of SAL (NaCl 0.9%) in the intraplantar surface of the mice right hind paw. Finally, all groups received their injection (PGE2 or SAL) at the same time of day (1 PM ± 1 hour, light cycle) starting and ending at 16-17 weeks old.

### 2.6 Delta Mechanical Nociceptive Threshold

The delta mechanical nociceptive threshold was obtained through electronic von-Frey (vF) apparatus (Insight, Riberão Preto, Brazil) adapted for mice(Deuis et al., 2017; Martinov et al., 2013). Briefly, the vF test was performed in a quiet, temperature-controlled room (21 ±1°C) and always at the same time of day (1 PM ± 1 hour, light cycle). Precisely, thirty minutes before the vF test, the mice were placed in an acrylic cage (12 × 20 × 17 cm) with wire grid floors for acclimation and the experimenters also remained in the room for appropriate environmental acclimation between the mice and experimenters as suggested by Martinov et al. (2013). To measure the hind paw mechanical nociceptive threshold, the experimenter applied constant pressure on the plantar surface of paws (right and left) until the paw-withdrawal threshold. The experimenter was blinded to PGE2-induced PH-ST protocol and to voluntary PA or sedentary behavior variables and partially blinded to diet variable because the mice of the HFD group were visibly larger than that of the SD group. The stimuli on the plantar surface were repeated (three-minimum or five-maximum times), not consecutively, until the measure could present three registrable measurements with equal values or with a difference less than 20% (0.2g) (Martinov et al., 2013). Those mice which did not complete these criteria of inclusion (*e.g.* five-maximum attempts) were excluded from the analysis.

The values of the mechanical nociceptive threshold were considered by the average of measurements performed at each time point assessment and the data were presented by the delta (Δ) of the mechanical nociceptive threshold. The delta was calculated by the difference between the average baseline values of the mechanical nociceptive threshold minus the average of the values of the mechanical nociceptive threshold at each respective time point assessment, such as before and after diets, before and after voluntary PA, after 3 hours of the first dose administration of PGE2-induced PH-ST protocol, and one and seven days after the end of PGE2-induced PH-ST protocol.

### 2.7 Voluntary Physical Activity

Voluntary PA behavior was evaluated through free RW access. Each RW had a magnetic indicator sensor that was connected to computer software to record summation of wheel revolution within five-minute intervals. The distance traveled was calculated based on RW interior diameter (9.2 cm) provided by Columbus Instruments (Columbus, OH, USA). Finally, we evaluated the mean of total daily distance traveled from 12 to 18 weeks old (6 weeks or 42 days in total) using a script routine in MATLAB® software (R12 version from MathWorks, Natick, MA, USA) and Microsoft Excel®.

### 2.8 Nucleus Accumbens Transcriptome Library Preparation

At 18 weeks old, the mice were euthanized by decapitation according to the American Veterinary Medical Association (American Veterinary Medical Association, 2013). The brain was immediately removed, snap-frozen in liquid nitrogen, and stored in a −80°C freezer. Bilaterally microdissected slices at 60 μm thickness of the NAc was processed using a cryostat (Leica® – Wetzlar, Germany) and were obtained from 1.96 mm up to 0.62 mm to bregma using the brain atlas (Franklin and Paxinos, 1996) and fixed on pre-prepared parafilm microscope slides. The parafilm microscope slices were a stain of Cresyl Violet 1% diluted in alcohol 50% (Sigma Aldrich) and with 40x optical microscopy, the NAc was identified and microdissected for transcriptome analysis.

Total RNA extraction of the NAc was obtained using Trizol® reagent (Invitrogen -Waltham, MA, USA) according to the manufacturer’s instructions. The RNA quality was confirmed by RNA 6000 Nano Assay for Agilent 2100 Bioanalyzer (Agilent Technologies – Stockport, United Kingdom). The RNA was quantified using a NanoVue Plus spectrophotometer (GE Healthcare Life Sciences – Pittsburgh, PA, USA) and a fluorometer (Qbuit®, Life Technologies – Carlsbad, CA, USA). The cDNA libraries were performed from 200 ng of extracted RNA using the TruSeq Stranded mRNA poliAAA (Illumina® – San Diego, CA, USA) according to manufacturer instructions. We performed 40 samples in two group-balanced lanes with 20 samples in each lane and 5 samples per group. The cDNA libraries were sequenced processed with an Illumina HiSeq® 2500 in High Output mode at the Macrogen NGS Service (Seoul, Republic of Korea), producing 101-bp paired-end reads. The first sequencing run produced a mean of 19,992,450 reads with a mean of 95.54% bases over Q30 per sample. The second run produced a mean of 18,090,251 reads with a mean of 96.94% bases over Q30 per sample. The sequence reads were aligned to the *Mus musculus* genome (GRCm38) from Ensembl (ensembl.org) using STAR 2.7.0 aligner (Dobin et al., 2013). Sequence read archive is accessible under number PRJNA564336 public repository at https://www.ncbi.nlm.nih.gov/.

### 2.9 Statistical Analyses

First, we performed the Shapiro-Wilk test to test for normality among the variables body mass, food intake, delta mechanical nociceptive threshold, and distance traveled. For all aforementioned variables, the analysis of variance with repeated measures (ANOVA-RM) was applied. For the adipose tissue variable, considering that voluntary PA may interfere on the amount of adipose tissue (epididymal and retroperitoneal) in both groups, a factorial ANOVA was applied. The Newman-Keuls post-hoc was applied when indicated. We also performed Grubb’s test to check outlier’s existence (*p*-value < 0.05) for all variables described above. Statistical testing was done using STATISTICA®10 software (StatSoft – Hamburg, Germany) and the data were presented by the mean and standard error of the mean (SEM) except for the body mass result, which was presented by the mean and standard deviation. Graphics were generated using GraphPad Prism 7 software (GraphPad Software – San Diego, CA. USA).

The interaction factors (diet *vs* RW *vs* PH-ST) of the differential gene expression analyses was performed using the DESeq2 R package library using a cutoff of an adjusted p-value of *P* < 0.05 (Love et al., 2014) and a Volcano R package (Blighe, 2019) was used to visualize the differential gene expression. The R script used is available in the supplementary material (Supple. A1). Functional annotation clustering was performed using the Database for Annotation, Visualization and Integrated Discovery (DAVID v6.8 https://david.ncifcrf.gov/) using DAVID EASE (Expression Analysis Systematic Explorer) score (*p* < .05), a modified Fisher’s exact test corrected for multiple hypothesis testing using Benjamini-Hochberg false discovery rates (FDR *p* < .05). The functional categories investigated included gene ontology (GO) of the biological processes.

## 3. Results

### 3.1 Body Mass, Caloric Intake and Adipose Tissue

The ANOVA-RM revealed a mean effect of diet (*F*_(1, 67)_ = 207.75, *p* <.001), time (*F*_(12, 804)_ = 745.86, *p* <.001) and an interaction effect of diet and time (*F*_(12, 804)_ = 234.67, *p* <.001) on the body mass throughout the experiment (**Figure 2**). Post-hoc analysis showed from the seventh to eighteenth weeks of age that the HFD group (24.27 ± .26g and 39.26 ± .62g, respectively) had significantly higher body mass when compared to an SD group (22.23 ± .27g, *p* = .004, and 26.24 ± .28g, *p* = .0001, respectively).

**Figure 2:**
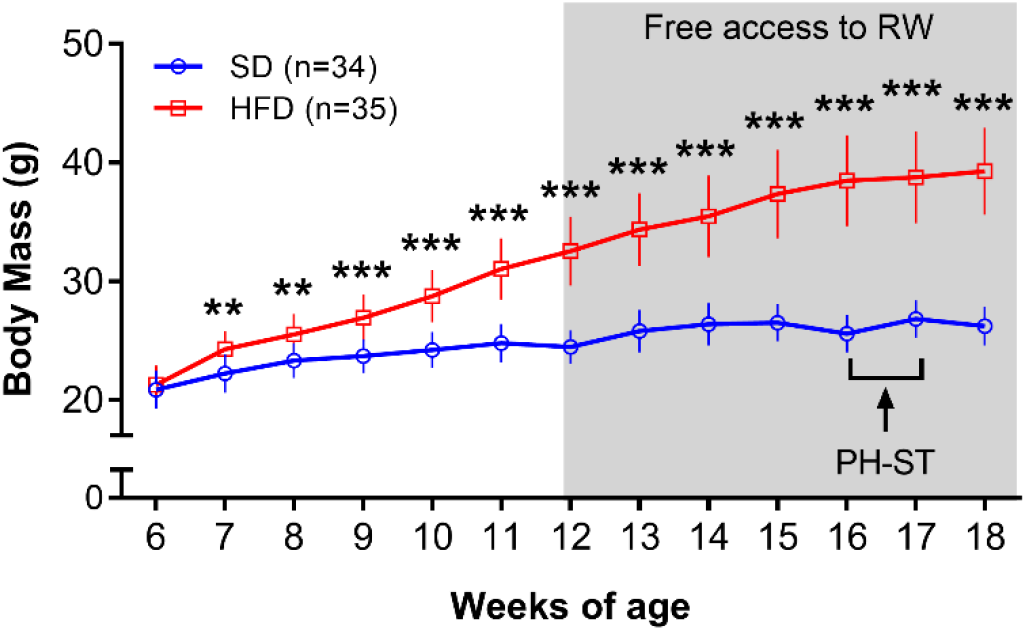
Body mass (grams) throughout the experiment. The ANOVA-RM and the post-hoc analyses revealed that the body mass was increased in the HFD group when compared to the SD group from the 7 into 19 weeks of age. Grey area represents the period of free access at a running wheel for voluntary physical activity. Data were presented by the mean and standard deviation. SD = standard diet, HFD = high-fat diet, RW = running wheel, PH-ST = persistent hyperalgesia short-term protocol. ** = *p* < .01 *** = *p* < .001

Moreover, during the period of free access to a RW for voluntary PA (12-18 weeks), the ANOVA-RM also showed a mean effect of diet (*F*_(1, 65)_ = 280.11, *p* <.001) and time (*F*_(6, 390)_ = 209.65, *p* <.001) but did not show a mean effect of PA (*F*_(1, 65)_ = 0.71, *p* = .40) on body mass. Moreover, the test showed an interaction effect of time and PA (*F*_(6, 390)_ = 4.27, *p* < .001) and an interaction effect of time and diet (*F*_(1,390)_ = 92.16, *p* < .0001) but, the test did not show an interaction effect of diet and PA (*F*_(1,65)_ = 0.17, *p* = .89) on body mass. The test also revealed an interaction effect of time, diet and PA (*F*_(6, 390)_ = 6.08, *p* <.001) on body mass during the period of free access to a RW for voluntary PA. Post-hoc analysis revealed that body mass were increased in the PA-HFD (32.25 ±2.72g) and SED-HFD (32.82 ±3.14g) groups when compared to PA-SD (24.59 ±1.41g, *p* = .0001 and *p* = .0001) and SED-SD (24.37 ±1.47g, *p* = .0001 and *p* = .0001) groups at the twelfth week of age into the eighteenth week of age: PA-HFD (40.24 ± 0.87g) and SED-HFD (38.22 ± 0.83g) when compare to PA-SD (26.74 ±0.35g, *p* = .0001 and *p* = .0001) and SED-SD (26.0 ± 0.44g, *p* = .0001 and *p* = .0001) (**Figure 3 A**). Post-hoc analysis also revealed that body mass had a significant increase at the pre and post moment to the free access to a RW within the SD (24.59 ± 0.34g *vs* 26.47 ± 0.35g, *p* = .0001) and the HFD (32.25 ± 0.64 *vs* 40.24 ±0.87, *p* < .0001) groups (**Figure 3 B**).

**Figure 3:**
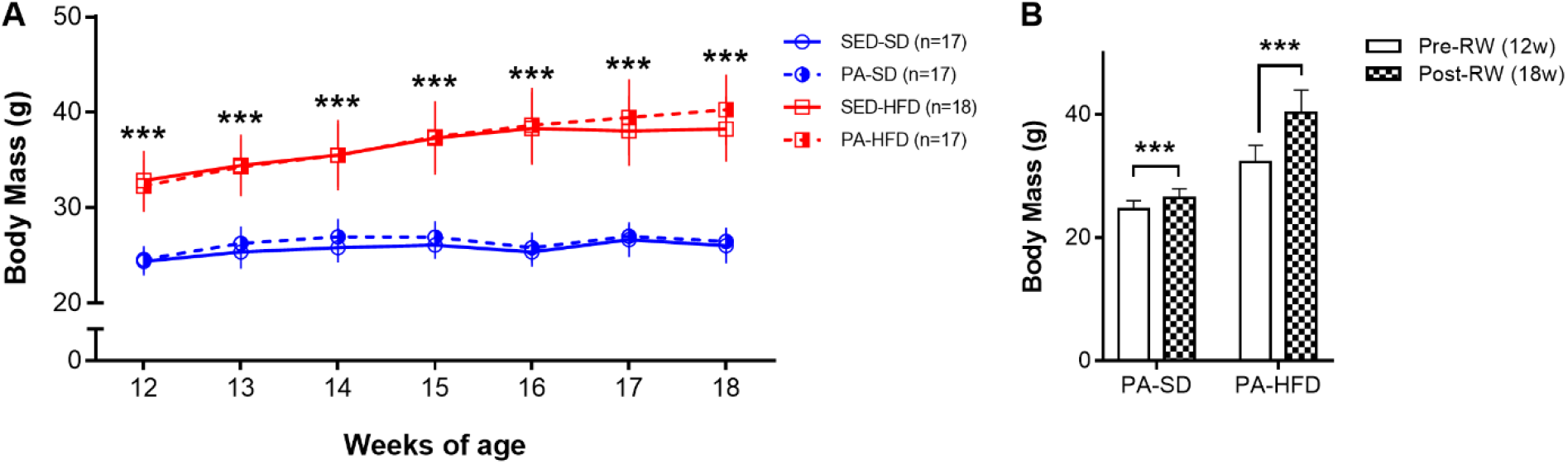
Body mass during the free access to running wheel for voluntary physical activity. In **A**, the analyses reveal that the SED-HFD and PA-HFD groups had significantly higher body mass when compared to SED-SD and PA-SD groups from twelfth until eighteenth week of age. In **B**, the ANOVA-RM and the post-hoc analyses reveal that the body mass had a significant increase at the pre-RW and post-RW moment within the SD and HFD groups (*** = *p* < .001). Data were presented by the mean and standard deviation. SED-SD = sedentary standard diet, PA-SD = physical activity standard diet, SED-HFD = sedentary high-fat diet, PA-HFD = physical activity high-fat diet groups. In **A**, *** = *p* < .001 for both SED-HFD and PA-HFD groups when compare to SED-SD and PA-SD groups.

The ANOVA-RM test revealed a mean effect of diet (*F*_(1, 67)_ = 14.51, *p* < .001), time (*F*_(11, 737)_ = 13.83, *p* < .001), and an interaction effect of diet and time (*F*_(11, 737)_ = 4.22, *p* < .001) on the caloric intake. Post-hoc analysis showed that the HFD group had a significant superior caloric intake when compared to SD group at the seventh (106.98 ± 1.46kcal/g *vs* 92.72 ± .1.79kcal/g, *p* = .001), eleventh (102.74 ± 1.97kcal/g *vs* 87.11 ± 2.17kcal/g, *p* = .0005), twelfth (101.19 ± 2.11kcal/g *vs* 86.92 ± 2.32kcal/g, *p* = .003), thirteenth (100.56 ± 1;80kcal/g *vs* 87.77 ± 2.5kcal/g, *p* = .012), fifteenth (100.52 ± 2.07kcal/g *vs* 99.59 ± 2.88kcal/g, *p* = .029) and sixteenth (107.11 ± 2.21kcal/g *vs* 94.78 ± 3.20kcal/g, *p* = .013) weeks of age old (**Figure 4**).

**Figure 4:**
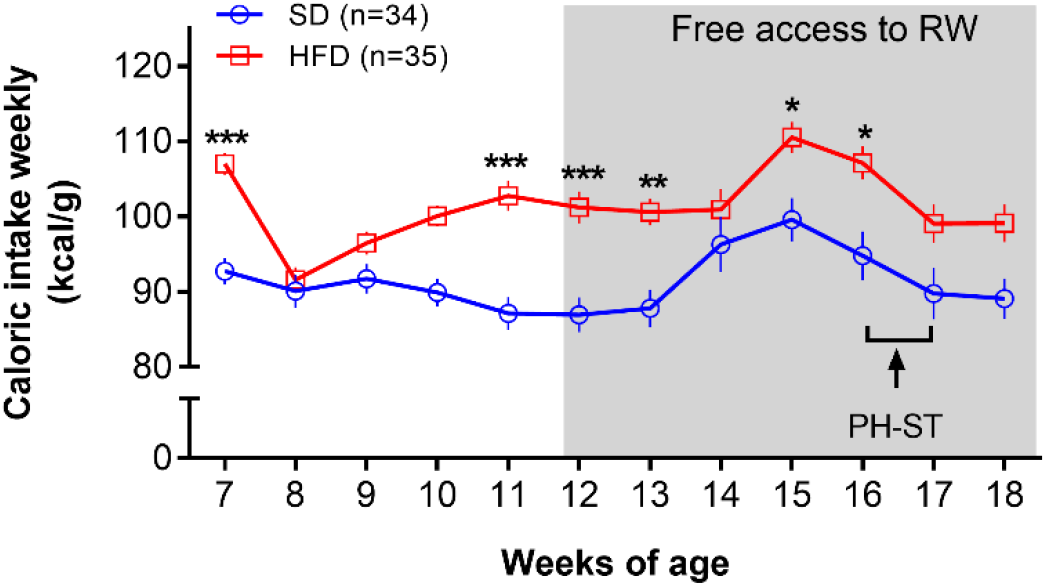
Caloric intake weekly. The ANOVA-RM and the post-hoc analyses revealed an increased caloric intake (kcal/g weekly) in the HFD group when compared to the SD group at the seventh, eleventh, twelfth, thirteenth, fifteenth and sixteenth weeks of age. The grey area represents the period of free access at a running wheel for voluntary physical activity. Data were presented by the mean and standard error of the mean. SD = standard diet, HFD = high-fat diet, RW = running wheel, PH-ST = persistent hyperalgesia short-term protocol. * = *p* < .05 ** = *p* < .01 *** = *p* < .001

The retroperitoneal and epididymal adipose tissues were higher in the HFD groups compared to SD groups. Factorial ANOVA revealed a mean effect of diet (*F*_(1, 65)_ = 617.15, *p* < .001) on retroperitoneal adipose tissue. However, the test did not reveal a mean effect of PA (*F*_(1, 65)_ = 0.75, *p* = .38) nor an interaction effect of diet and PA (*F*_(1, 65)_ = 0.45, *p* = .83) on retroperitoneal adipose tissue. Post-hoc analysis revealed retroperitoneal adipose tissue was significantly higher in the HFD groups (PA-HFD 1.48 ± 0.07g and SED-HFD 1.51 ± 0.06g) when compared to SD groups (PA-SD 0.20 ± 0.01g and SED-SD 0.26 ± 0.02g, *p* < .0001) (**Figure 5 A**). Factorial ANOVA also revealed a mean effect of diet (*F*_(1, 65)_ = 674.44, *p* < .001) on epididymal adipose tissue, however, the test did not reveal a mean effect of PA (*F*_(1, 65)_ = 0.92, *p* = .33). The test revealed a interaction effect of diet and PA (*F*_(1, 65)_ = 5.64, *p* = .02). Post-hoc analysis revealed epididymal adipose tissue was significantly higher in the SED-HFD (4.51 ± 0.82g) and PA-HFD (5.0 ± 0.87g) groups when compared to SED-SD (1.00 ± 0.18g, *p* < .001) and PA-SD (0.79 ± 0.13g, *p* < .001) groups (**Figure 5 B**). Post-hoc analysis also revealed epididymal adipose tissue of the PA-HFD group was significantly higher than the SED-HFD group (*p* = .021).

**Figure 5:**
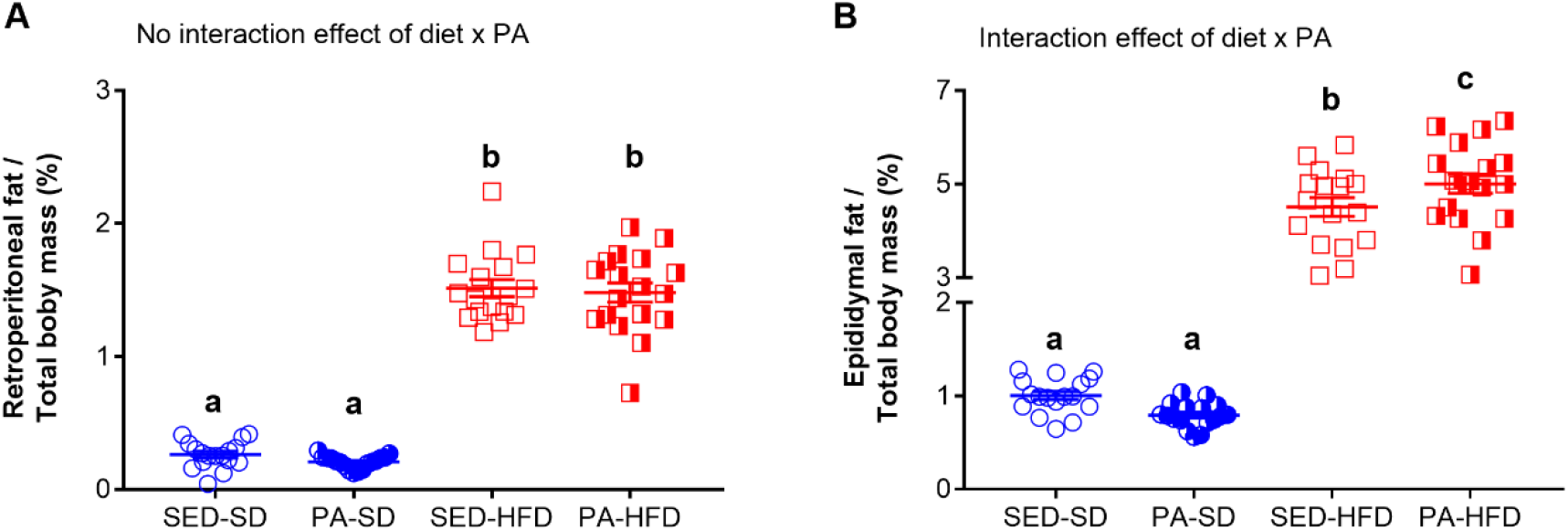
Retroperitoneal and epididymal adipose tissue. The factorial ANOVA and post-hoc analyses revealed that the retroperitoneal (**A**) and epididymal (**B**) adipose tissue were higher in the SED-HFD and PA-HFD groups when compared to the SED-SD and PA-SD groups. The test also revealed that the epididymal adipose tissue was higher in the PA-HFD group when compare to SED-HFD. SED-SD = sedentary standard diet group, PA-SD = physical activity standard diet group, SED-HFD = sedentary high-fat diet group and PA-HFD = physical activity high-fat diet group. n = 17-18 group. a *vs* b and c = *p* = .0001; b *vs* c = *p* = .021

### 3.2 Voluntary Physical Activity

Voluntary PA was decreased in the HFD group when compare to the SD group (**Figure 6**). In fact, the ANOVA-RM revealed a mean effect of diet (*F*_(1, 33)_ = 72.67, *p* < .001), time (*F*_(27, 891)_ = 46.26, *p* < .001), and an interaction effect of diet and time (*F*_(27, 891)_ = 11.58, *p* < .001) on distance traveled daily. Post-hoc analysis revealed that the HFD group showed a significant decrease on distance traveled from the sixth day (1.32 ± 0.29km) until the twenty-eighth day (2.51 ± 0.54km) when compared to the SD group at the sixth (4.57 ± 0.30km, *p* = .001) and twenty-eighth day (8.17 ± 0.56km, *p* = .0001).

**Figure 6:**
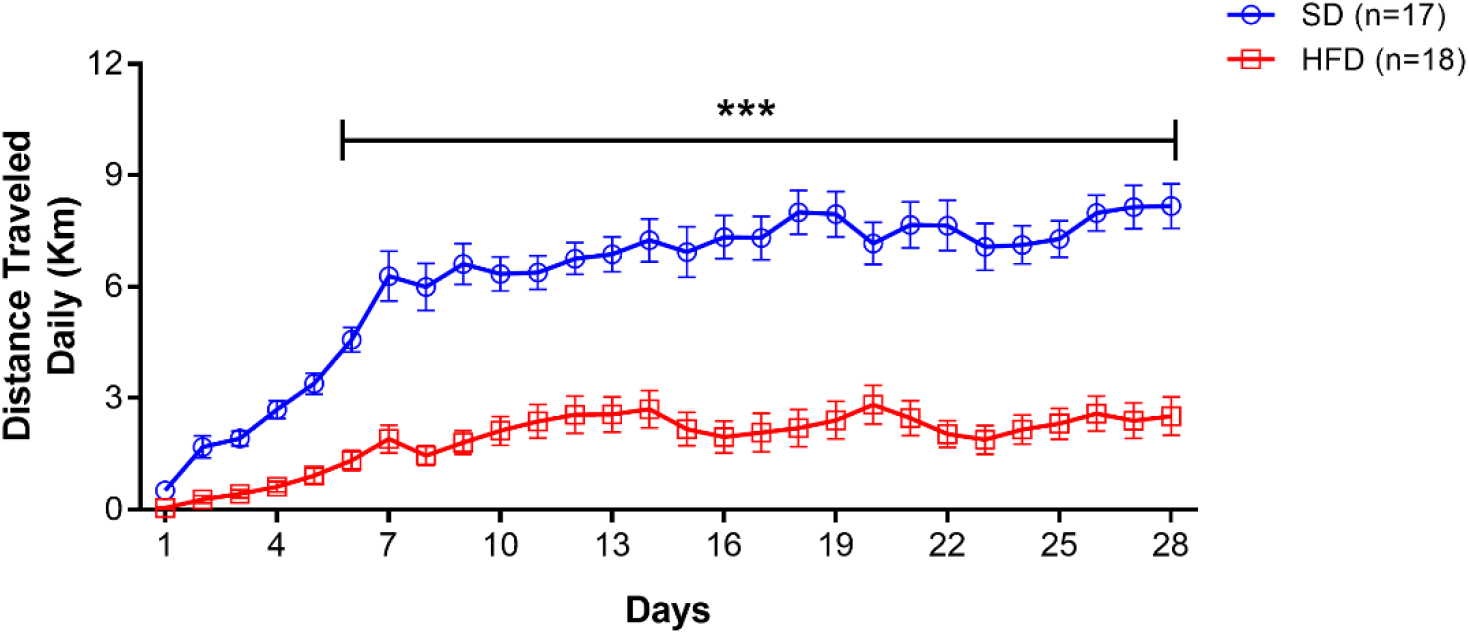
Voluntary physical activity (PA) measured by the distance traveled daily. The ANOVA-RM and the post-hoc analyses showed that the SD group had higher distance traveled when compared to the HFD group. Data were presented by the mean and standard error of the mean. SD = standard diet, HFD = high-fat diet. *** = *p* < .001.

### 3.3 High-fat Diet and Sedentary Behavior Promoted Chronic Pain Susceptibility

To test the effect of an HFD and sedentary behavior no promoting CP susceptibility, we first performed the ANOVA-RM test (diet *vs* PH-ST *vs* time). The test revealed a mean effect of PH-ST (*F*_(1, 30)_ = 38.87, *p* < .001) and time (*F*_(2, 60)_ = 61.08, *p* < .001), however the test did not reveal a mean effect of diet (*F*_(1, 30)_ = 2.32, *p* = .137) on PH susceptibility. On the other hand, the test revealed an interaction effect of diet and PH-ST (*F*_(1, 30)_ = 11.94, *p* = .001) and of diet and time (*F*_(2, 60)_ = 15.12, *p* < .001) on CP susceptibility. Moreover, the ANOVA-RM also revealed an interaction effect of PH-ST, diet, and time (*F*_(2, 60)_ = 12.52, *p* < .001) on CP susceptibility (**Figure 7**). Post-hoc analysis showed at one day before the beginning of PH-ST (−7 days **Figure 7**) that the delta mechanical threshold of the SED-HFD-PGE group (−1.37 ± .53g) was smaller when compared to SED-SD-PGE (0.25 ± .22g, *p* = .004) and SED-SD-SAL (−0.04 ±.34g, *p* = .016) groups, but not when compared to the SED-HFD-SAL (−0.82 ± .31g, *p* = .113) group. Further, at the first day after PH-ST post-hoc analysis showed the delta mechanical threshold was significantly higher in the SED-HFD-PGE group (3.79 ± .24g) when compared to SED-SD-PGE (2.74 ± .36, *p* = .028), SED-SD-SAL (0.91 ± .34, *p* = .0001), and SED-HFD-SAL (0.76 ± .19, *p* = .0001) groups. The delta mechanical threshold also was significantly higher in the SED-SD-PGE group when compare to SED-SED-SAL (*p* = .0006) and SED-HFD-SAL (*p* = .0004) groups. Finally, at the seventh day after PH-ST, post-hoc analysis showed that the delta mechanical threshold was significantly higher only in the SED-HFD-PGE (3.85 ± .20) group when compare to SED-SD-PGE (0.48 ± .19g, *p* = .0001), SED-HFD-SAL (0.70 ± .46g, *p* = .0001), and SED-SD-SAL (1.05 ± .36g, *p* = .0001) groups (**Figure 7**) revealing the effect of an HFD and sedentary behavior were able to promote CP susceptibility.

**Figure 7:**
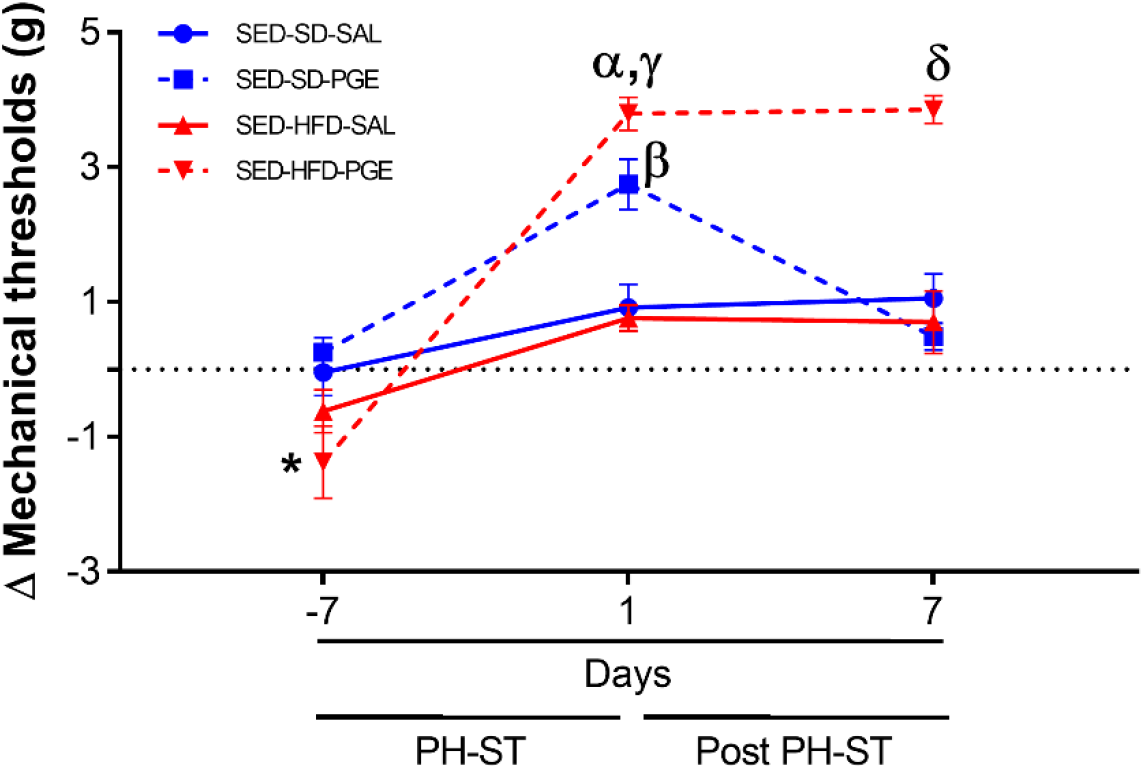
Chronic pain susceptibility promoted by HFD and sedentary behavior. The ANOVA-RM and post-hoc analyses revealed on the seventh day after PH-ST that the delta mechanical threshold was significantly higher in the SED-HFD-PGE group when compare to SED-SD-PGE, SED-HFD-SAL, and SED-SD-SAL groups. SED-SD-SAL and SED-HFD-SAL (n=8), SED-SD-PGE and SED-HFD-PGE (n=9). Data are presented by means and standard error of the mean. * = *p* < .05. SED-HFD-PGE ≠ SED-SD-PGE, SED-HFD-SAL, and SED-SD-SAL one day before PH-ST. α = *p* < .001. SED-HFD-PGE ≠ SED-SD-SAL and SED-HFD-SAL one day after PH-ST. γ = *p* = .02. SED-HFD-PGE ≠ SED-SD-PGE one day after PH-ST. β = *p* < .001 SED-SD-PGE ≠ SED-HFD-SAL and SED-SD-SAL one day after PH-ST. δ = *p* < .001 SED-HFD-PGE ≠ SED-SD-PGE, SED-HFD-SAL and SED-SD-SAL seven days after PH-ST.

### 3.4 Voluntary Physical Activity Prevented Chronic Pain Susceptibility

To test the effect of PA to prevent CP susceptibility, we performed the same analyses described in section 3.3, albeit within all physical activity groups. Thus, the ANOVA-RM revealed a mean effect of PH-ST (*F*_(1, 31)_ = 19.47, *p* < .001), diet (*F*_(1, 31)_ = 5.23, *p* = .02), and time (*F*_(2, 62)_ = 60.28, *p* < .001) on the delta mechanical threshold. The test also revealed an interaction effect of time and diet (*F*_(2, 62)_ = 16.19, *p* < .001) but the test did not reveal an interaction effect of PH-ST and time (*F*_(1, 31)_ = 1.90, *p* = .177) on the delta mechanical threshold. Moreover, the ANOVA-RM revealed an interaction effect of PH-ST, time, and diet (*F*_(2, 62)_ = 4.32, *p* = .017) on the delta mechanical threshold (**Figure 8**). Nonetheless, differently among sedentary groups, the post-hoc analysis showed at one day before the beginning of PH-ST that the delta mechanical threshold of the PA-HFD-PGE (−1.02 ± .46g) group was not significantly different when to compare to all groups: PA-SD-PGE (−0.14 ± .33g, *p* = .17), PA-HFD-SAL (−0.76 ± .43g, *p* = .59), and PA-SD-SAL (0.32 ± .18g, *p* = .056) groups. Further, at one day after PH-ST, post-hoc analysis showed the delta mechanical threshold was significantly higher in the PA-HFD-PGE group (3.95 ± .31g) when compared to the PA-SD-PGE (2.6 ± .37g, *p* = .007), PA-HFD-SAL (2.06 ± .44g, *p* = .001), and PA-SD-SAL (0.15 ± .26g, *p* = .0001) groups. Unexpectedly, the delta mechanical threshold was also significantly higher in the PA-HFD-SAL group when compared to the PA-SD-SAL group (*p* = .002) but was significantly smaller when compared to the PA-HFD-PGE group (*p* = .001). Finally, post-hoc analysis showed that the delta mechanical threshold was significantly decreased at seven days after the PH-ST protocol in the PA-HFD-PGE group (2.5 ± .46g) when compared to itself one day after the PH-ST (3.95 ± 31g, *p* = .008). Moreover, the same delta mechanical threshold of the PA-HFD-PGE group (2.5 ± .46g) at seven days was similar when compare to the PA-SD-PGE (2.6 ± .37g, *p* = .84) and to the PA-HFD-SAL (2.06 ±.44g, *p* = .37) groups also at one day after the PH-ST (**Figure 8**). This suggests the effect of PA in preventing CP susceptibility development promoted by HFD and sedentary behavior. The analyses also showed that the delta mechanical threshold was significantly higher at seven days after the PH-ST protocol in the PA-HFD-PGE group when compared to the PA-SD-PGE (0.57 ± 0.21g, *p* = .0006), PA-HFD-SAL (0.33 ± 0.19g, *p* = .0004), and PA-SD-SAL (0.55 ± 0.21g, *p* = .001) groups.

**Figure 8:**
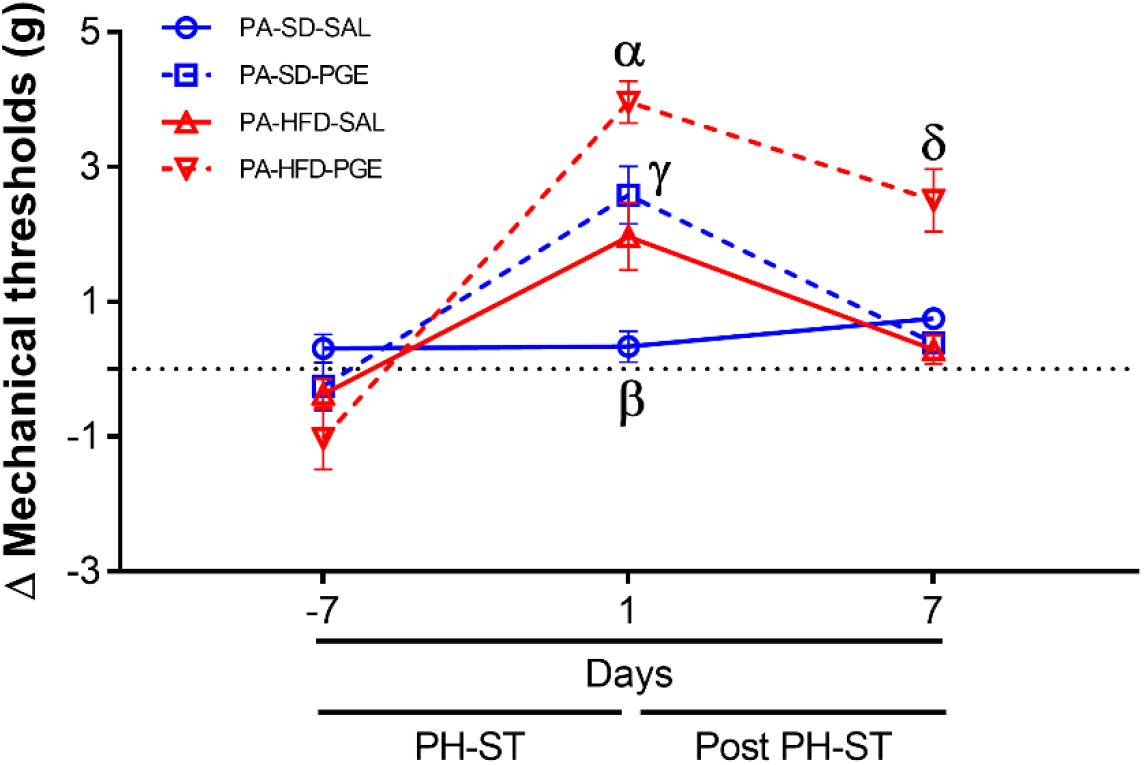
Physical activity (PA) prevented chronic pain susceptibility. The ANOVA-RM and post-hoc analyses revealed at the seventh day after the PH-ST that the delta mechanical threshold was significantly decreased in the PA-HFD-PGE group when compared to the PA-HFD-PGE group at one day after PH-ST and was similar when compared to PA-SD-PGE and PA-HFD-SAL at one day after PH-ST. PA-HFD-PGE, PA-HFD-SAL, and PA-SD-PGE (n=9). PA-SD-SAL (n=8). Data are presented by means and SE. α = *p* < .001. PA-HFD-PGE ≠ PA-SD-PGE, PA-HFD-SAL, and PA-SD-SAL one day after PH-ST. γ = *p* = .53. PA-HFD-SAL = PA-SD-PGE one day after PH-ST. β = *p* < .01. PA-SD-SAL ≠ PA-HFD-SAL one day after PH-ST. δ ≠ α = *p* = .008. PA-HFD-PGE seven days after the PH-ST ≠ PA-HFD-PGE one day after the PH-ST. δ = γ *=* PA-HFD-PGE seven days after PH-ST = PA-SD-PGE (*p* = .37) and PA-HFD-SAL (*p* = .84) one day after PH-ST. δ = *p* ≤ .001. PA-HFD-PGE ≠ PA-SD-PGE, PA-HFD-SAL and PA-SD-SAL seven days after the PH-ST.

### 3.5 Differential Gene Expression in the Nucleus Accumbens – Transcriptome Analysis

To investigate the differential gene expression in the NAc related to CP susceptibility promoted by an HFD and sedentary behavior and prevented through voluntary PA, we first compared the differential gene expression of the SED-HFD-PGE, a CP susceptibility group, (**Figure 7**) and PA-HFD-PGE, a prevented CP susceptibility group (**Figure 8**). This analysis revealed that 2,204 genes were differentially expressed in the NAc. The analysis also revealed 1,098 up-regulated and 1,106 down-regulated genes in the PA-HFD-PGE group thereby identifying genes involved in preventing and promoting CP susceptibility, respectively (**Figure 9** and Supplementary Table S1).

**Figure 9:**
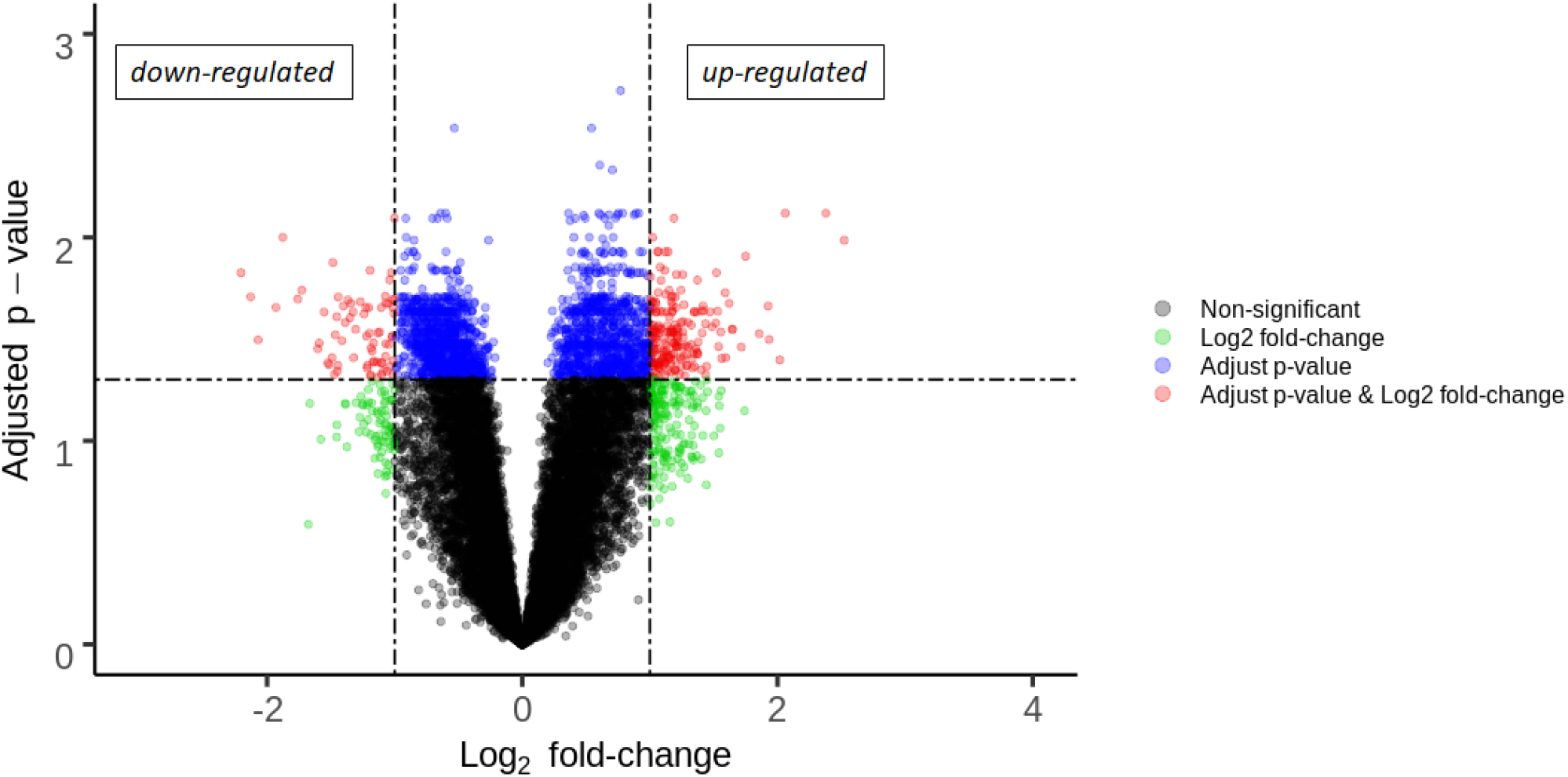
Volcano plot of the differential gene expression analyses in the nucleus accumbens (NAc). Analysis of NAc through RNA-Seq revealed 2,204 differential gene expression with 1,098 up-regulated and 1,106 down-regulated genes implicated in the prevention and promoting CP respectively. Adjusted p-value *p* = 0.05

According to DAVID gene ontology enrichment results, these genes were implicated in 41 biological processes (EASE *p* < .001 and FDR *p* < .05): 23 biological processes were enriched when considering up-regulated genes and 18 were enriched when considering down-regulated genes. One biological process was co-up-down-regulated (GO:0006810 transport). The 10 lowest *p*-values from the up and down-regulated genes related to the biological processes are presented in **Figure 10** and the gene list is presented in **Tables 3** and **4**. The total list of biological processes and differential gene expression in the NAc implicated in CP susceptibility promoted by an HFD and sedentary behavior in the SED-HFD-PGE group and prevented by voluntary PA (PA-HFD-PGE group) were described in the supplementary material (Table S2 and S3).

**Figure 10:**
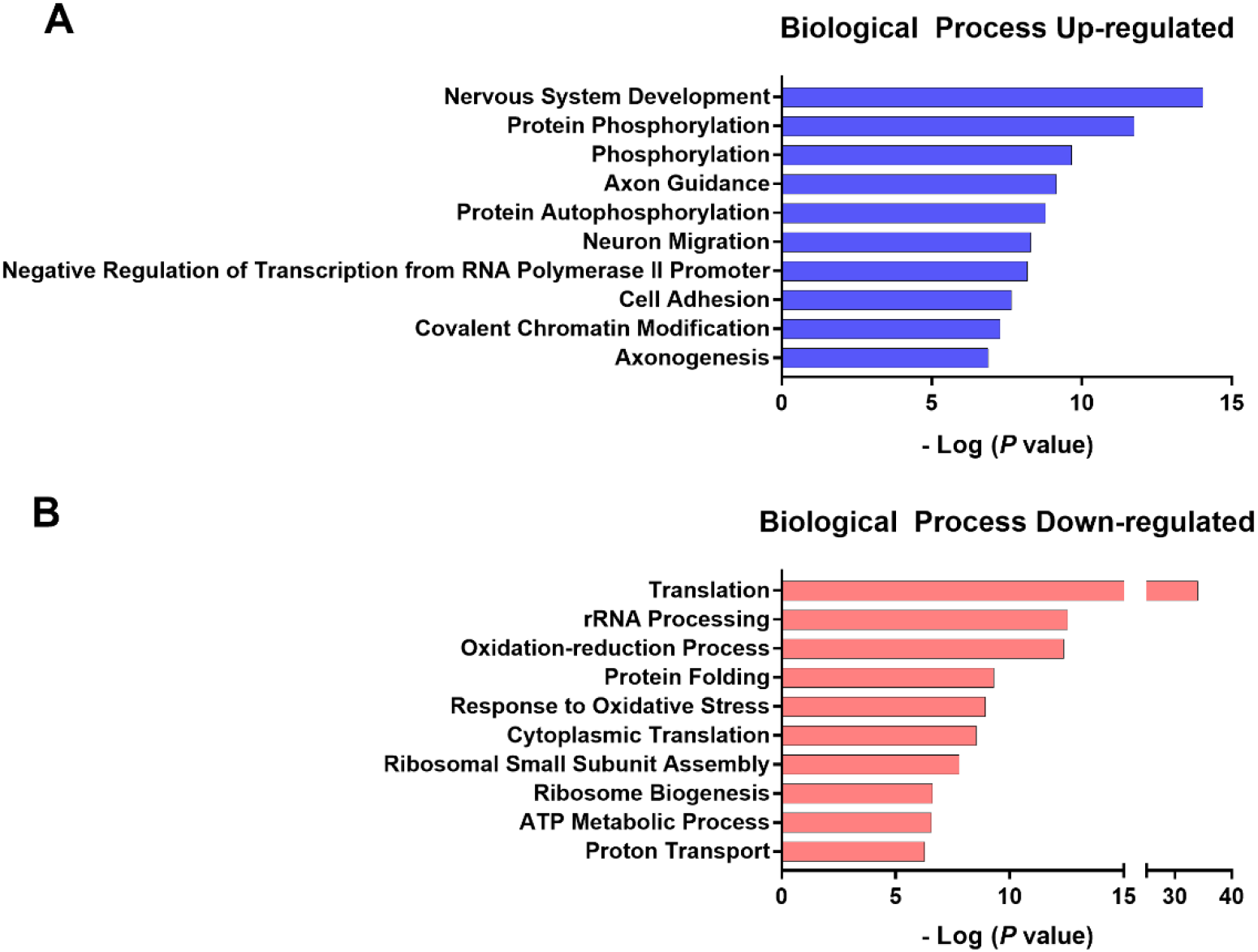
The ten lowest *p*-value up and down-regulated of biological processes. The ten lowest normalized *p*-value of biological processes up- (**A**) and down- (**B**) regulated genes related to chronic pain (CP) susceptibility induced by HFD and sedentary behavior and prevented by PA.

**Table 3:**
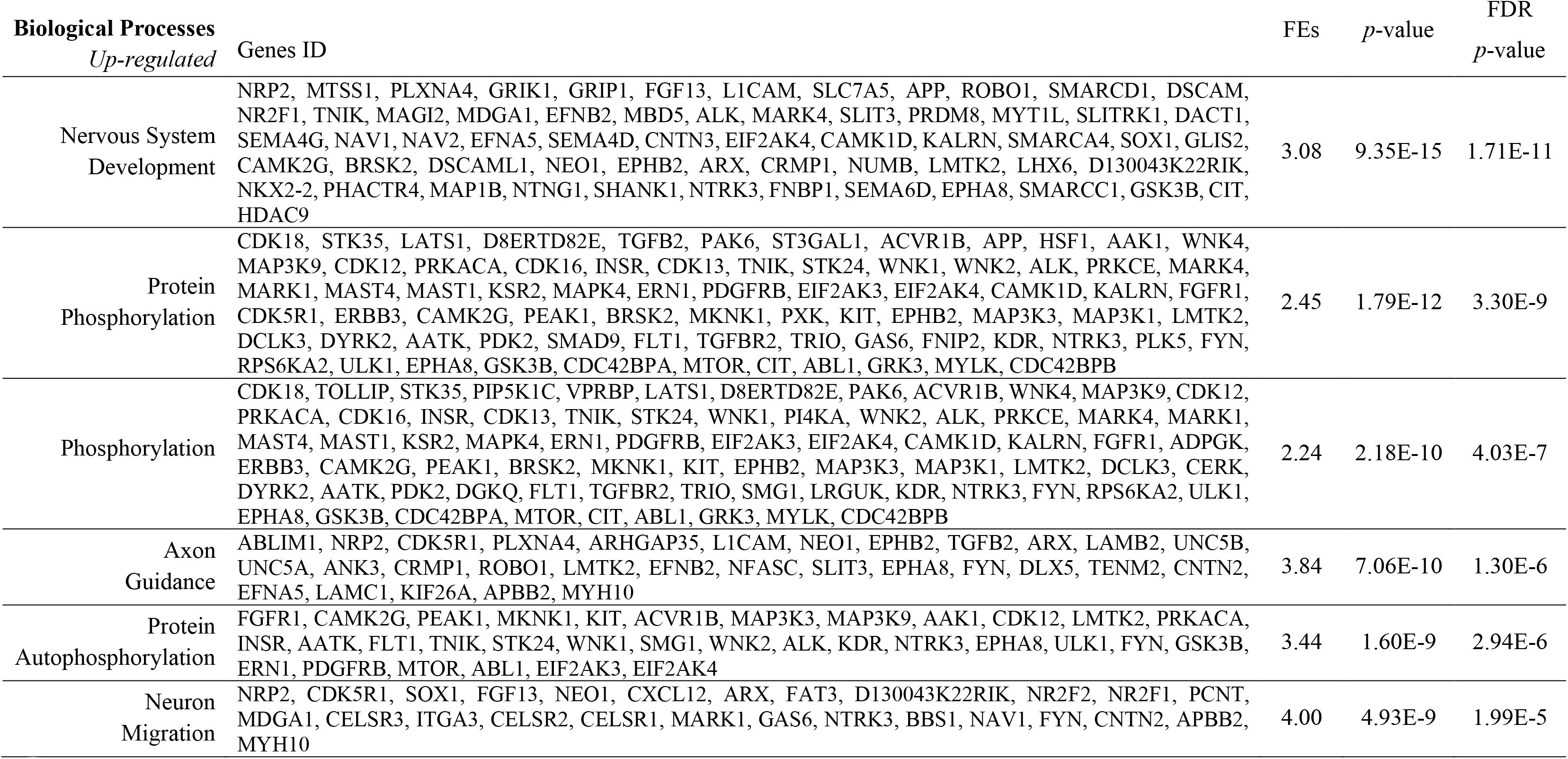
List of the biological processes and genes IDs of the six lowest *p*-value up-regulated genes enriched. FEs= fold enrichment score, FDR = false discovery rate

**Table 4:** List of the biological processes and genes IDs of the six lowest *p*-value down-regulated genes enriched. FEs= fold enrichment score, FDR = false discovery rate

| Biological Processes | Genes ID | FEs | *p*-value | FDR<br>*p*-value |
| --- | --- | --- | --- | --- |
| *Down-regulated* |  |  |  |  |
| Translation | RPL18, RPL17, RPL36A, RPL19, MRPL41, RPL14, RPL13, RPL15, RPS27L, RPL22L1, MRPL36, RPL10, FAU, RPL11, RPL12, RPS27A, 2810006K23RIK, RPL35A, MRPL52, MRPL51, RPS18, RPS19, SLC25A34, RPS16, RPS17, RPS14, RARS, RPS15, MRPL49, RPS12, GATB, MRPS17, EEF1B2, MRPS14, FARS2, MTIF3, QRSL1, RPS26, RPS27, RPS28, RPL7, RPS29, RPL6, RPL9, EIF3K, RPL3, MRPL55, EIF3L, EIF3I, RPL5, RPL10A, RPL7A, RPS20, RPS21, RPS23, EIF3M, RPS24, RPSA, MRPS24, RPL23A, MRPS21, DENR, RPS6, MRRF, RPS5, RPS8, RPS3A1, RPS7, SLC25A14, RPL18A, RPL37A, RPS3, MRPS5, RPS4X, RPL41, EIF4A1, FARSB, NHP2, RPL27A, RPL35, RPL36, RPS15A, RPL37, RPL38, RPL39, MRPL20, RACK1, MRPL11, RPL30, MRPL15, RPL32, RPL31, RPL34, MRPL18, RPS27RT, AIMP1, AIMP2, RPL26, RPL27, MRPL30, RPL29, MRPL24, MRPL23, MRPL21, RPL23, RPL22, RPL13A, RPL21 | 5.50 | 9.17E-50 | 1.62E-46 |
| rRNA processing | RPL14, LSM6, FCF1, SBDS, IMP3, RPS28, RPL7, RPL5, RPL11, MTERF4, FTSJ3, RPS24, RPL35A, EXOSC9, EXOSC7, EMG1, RPL26, GTF2H5, RPS6, NOP10, RNMTL1, MRM2, RPS7, RPS19, PIH1D1, RPS16, RPS17, RPS15, POP5, MPHOSPH6, NHP2 | 5.02 | 2.96E-13 | 5.24E-10 |
| Oxidation-Reduction<br>Process | LDHB, LDHA, GMPR2, PRDX5, RPE65, PRDX2, UQCRQ, PRDX1, FDFT1, NDUFS6, GPX1, HMOX2, UQCR10, MSRA, NDUFS5, UQCR11, NDUFS4, IDH3G, HMOX1, GPX4, NDUFS8, NDUFS3, SUOX, NDUFB11, NDUFC2, CYB5A, DECR1, CBR3, COQ7, CDO1, DHRS7B, DDO, NDUFA12, NDUFA11, DHRS1, DHRS4, PRDX6, MARC2, UQCRH, UQCRB, PRODH, HSD17B11, NDUFB3, HSD17B10, NDUFB5, NDUFB6, NDUFB7, TXN2, NDUFB9, ADH5, TXN1, SESN1, HADHA, CYB561D2, AKR1A1, FMO1, FAM213A, NDUFA4, NDUFA5, NDUFA2, NDUFA3, NDUFA9, NDUFA6, NDUFA7, FAM213B, SOD1, NDUFA1, MSRB2, BLVRA, NDUFV3, SDHB, TXNDC12, AKR1B3, HSDL2, BLVRB, NDUFV2, PHGDH, CRYM, DCXR, MGST1 | 2.41 | 4.36E-13 | 7.72E-10 |
| Protein Folding | FKBP8, GRPEL1, PDIA3, TXN2, FKBP4, PDIA6, TXN1, CCT3, HSCB, CDC37, PPIL3, PDRG1, HSPE1, TUBA1B, HSPA8, TCP1, DNLZ, PPIE, PFDN1, PPIH, HSP90B1, CCT5, PPIB, PPIA, PFDN5, TBCC, AHSA1 | 4.30 | 4.90E-10 | 8.68E-7 |
| Response to<br>Oxidative Stress | NDUFB4, ATOX1, PRDX5, PINK1, PRDX2, PRDX1, RPS3, PSMB5, GPX1, HMOX2, MSRA, APOE, GPX4, HMOX1, TOR1A, NDUFS8, ERCC1, MT3, SELK, NDUFA6, CST3, SOD1, COQ7, MSRB2, PARK7, NDUFA12, PEBP1 | 4.14 | 1.17E-9 | 2.07E-6 |
| Cytoplasmic<br>Translation | RPL35A, RPL15, RPL26, RPL36, RPL22L1, RPL29, RPL7, RPL31, RPL6, RPL22, RPL9, RPLP0, RPLP1, FTSJ1 | 8.41 | 2.97E-9 | 5.26E-6 |

Furthermore, we compared the differential gene expression in the NAc of the SED-HFD-PGE, a CP susceptibility group, versus the SED-SD-PGE, a non-CP susceptibility group (**Figure 7**). We also compared the PA-HFD-PGE, a prevented CP susceptibility group and versus the PA-SD-PGE, a non-CP group (**Figure 8**). Both analyses did not reveal any differential gene expression in the NAc. Confirming our hypothesis that CP susceptibility was promoted when an HFD was combined with sedentary behavior and that voluntary PA *per se* prevented CP susceptibility even when the mice PA group were fed an HFD.

## 4. Discussion

We presented here the first report of differential gene expression in the mice NAc implicated in CP susceptibility promoted by HFD and sedentary behavior but which in turn was prevented by voluntary PA, even when the mice was fed an HFD. We first showed a PGE-induced PH-ST protocol, which was not enough to induce CP on mice fed an SD when they were sedentary, it was enough to induce CP in the SED-HFD group, demonstrating that the HFD and sedentary behavior promoted CP susceptibility (**Figure 7**). Second, we our results showed that voluntary PA group (PA-HFD-PGE) was prevented to CP susceptibility (**Figure 8**). These results confirmed our hypothesis that an HFD and sedentary behavior lead to CP susceptibility, while voluntary PA can prevent it. Previous studies have documented the role of HFD in nociception modulation and CP (Cooper et al., 2018b; Liang et al., 2019; Rossi et al., 2013; Song et al., 2017), however, our report is the first to investigate the implication of differential gene expression and the biological processes in the NAc involved to CP susceptibility. To the best of our knowledge, this study is also the first to report that biological processes implicated to neurodegeneration and neuroplasticity in the NAc promoted and prevented CP susceptibility, respectively.

The results of interaction factors analysis from the transcriptome data of the NAc samples (**Figure 9**), revealed that 1,098 up-regulated and 1,106 down-regulated genes were implicated in preventing and promoting susceptibility to CP. Although our study has been the first to employ a transcriptomic approach of the NAc to investigate susceptibility to CP promoted by an HFD-induce obesity and sedentary behavior, other studies have shown that differential gene expression in the NAc was related to CP and other comorbidities. For instance, in an animal study applying transcriptomic to the NAc, mPFC, and PAG by Descalzi et al. (2017) showed that neuropathic pain promoted gene expression changes in these brain areas and those changes were involved in stress and depression comorbidities. Curiously, some biological processes related to neuropathic pain found in the Descalzi et al. (2017) study were similarly related to CP susceptibility found in our study, such as phosphorylation and regulation of transcription from RNA polymerase II promoter (**Figure 10 A**). Although Descalzi et al. (2017) study did not investigate the effect of an HFD and voluntary PA, we reported here that these variables play a crucial role in CP susceptibility and other comorbidities such as obesity.

Through gene ontology enrichment analysis of differential gene expression in the NAc (**Figure 10 B** and Table S2, S3) 41 biological processes were related to preventing and promoting of CP susceptibility. We identified 23 biological processes enriched when considering up-regulated genes in the SED-HFD-PGE group implicated with CP susceptibility promoted by a HFD and sedentary behavior, such as, translation (GO:0006412), response to oxidative stress (GO:0006979), ATP metabolic process (GO:0046034), response to oxidative stress (GO:0006979) proton transport (GO:0015992), ATP synthesis couple proton transport (GO:0015986). The gene ontology enrichment analysis also revealed 18 biological processes enriched when considering up-regulated genes in the PA-HFD-PGE group implicated in the prevention of CP susceptibility (**Figure 10 A** and Table S2, S3), such as nervous system development (GO:0007399), neuron migration (GO:0001764), axonogenesis (GO:0007409), protein autophosphorylation (GO:0006468), cell adhesion (GO:0007155), actin cytoskeleton organization (GO:0030036), learning (GO:0007612). Taken together, these data reported here suggesgt that biological processes involved in phosphorylation, as reported by Descalzi et al. (2017), metabolic and mitochondrial stress in the NAc may be associated with an increased susceptibility to CP, whereas biological processes involved in neuroplasticity and axonogenesis in the NAc limits susceptibility to CP, at least when caused by an HFD and sedentary behavior. In fact, we were the first to report that biological processes involved in neuroplasticity, at least within an HFD-induce obesity and sedentary behavior context, may prevent the CP susceptibility. Whereas several studies reported that neuroplasticity in the NAc is underlying CP in humans and animals (Apkarian et al., 2011; DosSantos et al., 2017; Kuner and Flor, 2016; Woolf, 2000), our results may reflect an neurodegeneration process in the NAc caused by HFD and sedentary behavior supporting the CP susceptibility, and secondarily, a neuroplasticity counter-regulatory mechanism orchestrated by the voluntary PA. Indeed, PA and physical exercise is already known to produce neuroplasticity (Chen et al., 2017; Fernandes et al., 2017; Lourenco et al., 2019).

We also reported that the SED-HFD-PGE group was with CP susceptibility when compared to the SED-SD-PGE or to the SED-HFD-SAL groups seven days after the end of the PGE-induced PH-ST protocol, confirming that CP susceptibility was promoted by an HFD and sedentary behavior (**Figure 7**). These findings support a study by Song et al. (2017) when they found an increase of pain behaviors promoted by an HFD regardless of weight gain. Although the authors found an increased macrophage density in the dorsal root ganglia and increased pain behaviors (Song et al., 2017), they did not investigate any potential effect of an HFD and sedentary behavior in the NAc, a key structure in the CP susceptibility (Dias et al., 2015; Schwartz et al., 2017; Zhang et al., 2019).

Furthermore, we reported here that the voluntary PA prevented CP susceptibility promoted by an HFD and sedentary behavior. In fact, the result revealed that the PA-HFD-PGE group presented a significant decrease in the delta mechanical threshold at seven days after the PH-ST protocol when compared to one day after the PH-ST protocol (**Figure 8**). The results at seven days after the PH-ST protocol also showed that the PA-HFD-PGE group was with the similar delta mechanical threshold of non-CP groups (PA-SD-PGE and PA-HFD-SAL) at one day after the PH-ST protocol. Despite the PA-HFD-PGE group having a higher mechanical threshold when compared to PA-SD-PGE, PA-HFD-SAL, or PA-SD-SAL groups at seven days after the PH-ST protocol, taken together, these results support the hypothesis that voluntary PA prevents CP susceptibility promoted by an HFD and sedentary behavior. According to our knowledge, those results regarding the effect of voluntary PA in preventing CP susceptibility promoted by an HFD and sedentary behavior was not previously described in the literature.

Despite not finding a decreased body mass in the voluntary PA groups (**Figure 2**), a meta-analyses study reported that weight mass reduction promoted by PA or other interventions such as caloric diet-restriction can contribute to pain reduction in humans with chronic musculoskeletal pain (Cooper et al., 2018a). The contrast between these results suggests that the effect of voluntary PA in preventing CP susceptibility in mice or pain reduction in humans with CP may be mediated by differential gene expression related to neuroplasticity in the NAc as we report here. Moreover, although Sluka et al (2013) and Leung et al (2016) have described the effects of regular PA to prevent CP development in animals models, our study is the first to report the interaction between diet (SD or HFD), sedentary behavior (PA or SED), and describe the biological processes in the NAc underlying CP susceptibility. Thus, taken together, our results offer novel insights into molecular mechanisms and biological processes underlying the development of CP susceptibility within the individuals with obesity and sedentary lifestyles.

As expected and widely reported in the literature (Kleinert et al., 2018; Montgomery et al., 2013; Williams et al., 2014; Yang et al., 2014), after 12 weeks on a HFD the body mass higher in the HFD group compared to the SD group (**Figure 2**). However, our result showed that the 6 weeks of voluntary PA was not enough to reduce body mass in the PA-HFD when compare to the SED-HFD group or in the PA-SD when compared to the SED-SD groups (**Figure 2** and **Figure 3**). As our aim in this study was to investigate the effect of an HFD, sedentary behavior and voluntary PA in promoting or preventing CP susceptibility, the body mass stability throughout our study (**Figure 3 A** and **B**), even after the PA period, increased our confidence that CP susceptibility results are not related to the body mass change, as reported by Song et al (2017). Moreover, body mass result reported here is in accordance with a Rocha-Rodrigues et al. (2018) study that revealed no significant difference in rats body mass after 9 weeks on an HFD when the animals were submitted in 8 weeks of running protocol (treadmill exercise or voluntary PA). It seems whether mice start voluntary PA belatedly, (*e.g.* 6 weeks in our study after initiating an HFD), the activity tends to produce no effect on the total body mass or in the amount of adipose tissue.

At the same time, caloric intake was higher in the HFD versus the SD group (**Figure 4**) and this result is in accordance with the literature (Hu et al., 2018a; Kleinert et al., 2018; Yang et al., 2014). The high energy density found in the HFD (5.439 kcal/g) compared to the SD (3.080 kcal/g), particularly from fat (58.2% *vs* 11.7%, respectively), support the caloric intake higher in the HFD group and it is in accordance to a recent study where the authors suggested that the amount of fat from an HFD, but not protein or carbohydrate, is the cause of adiposity increases in mice (Hu et al., 2018a). Further, we also found an increase of adipose tissue in the HFD groups when compared to SD groups for retroperitoneal (**Figure 5 A**) and epididymal adipose tissue (**Figure 5 B**). Nonetheless, we found an unexpected interaction effect of voluntary PA and HFD in the epididymal tissue. We found a significantly higher amount of epididymal adipose tissue in the PA-HFD group when compared to the SD-HFD group (**Figure 5 B**). Unlike our findings, a previous study showed that forced physical exercise on a treadmill was associated with epididymal adipose tissue decrease when mice were fed an SD (Castellani et al., 2014). However, conversely, Rocha-Rodrigues et al. (2018) reported no difference in epididymal adipose tissue when mice were fed an HFD nine weeks before the start of a voluntary PA protocol.

Nevertheless, the aim of our study was not to investigate changes in adipose tissue related to CP susceptibility, it appears inflammatory profile changes of adipose tissue may play a role in CP susceptibility at a peripherical level. Hu et al. (2018b) demonstrated that leptin mediated formalin-induced nociception in mice, whereas Li et al. (2013) demonstrated that intrathecal leptin administration alleviated neuropathic pain induced by sciatic chronic constriction. Thus, our results raise other interesting questions that remain uninvestigated. For instance, whether gene expression changes found in the NAc are caused by inflammatory profile changes at the peripherical level.

Another interesting finding in our study that remains uninvestigated is related to the effect of an HFD in the motivation for voluntary PA (**Figure 6**) under the perspective of gene expression changes in the NAc. Other researchers have already shown a decrease of distance traveled in voluntary PA when mice were fed an HFD and high-sugar diet or a typical western diet compared to mice fed an SD (Bjursell et al., 2008; Funkat et al., 2004; Vellers et al., 2017). However, few studies investigate these questions from a NAc-centric perspective, a key structure for motivation (Salamone et al., 2015; Salamone and Correa, 2012). In one study, the authors reported basal ganglia dysfunction in striatal D2R contribute to physical inactivity in obesity and the physical inactivity was not related to an increase of body mass or obesity, suggesting that physical inactivity is not caused by obesity (Friend et al., 2017), however, the differential gene expression in the NAc related to physical inactivity is unknown, confirming the lack of understanding regarding the effect of an HFD in the NAc gene expression and motivation for voluntary PA.

## 5. Conclusion

To conclude, first, we reported here that an HFD and sedentary behavior promoted CP susceptibility in mice, whereas voluntary PA prevented it. Second, differential gene expression and gene ontology enrichment in the NAc analysis revealed that biological processes implicated to neurodegeneration were involved in promoting CP susceptibility. We also concluded that biological processes implicated in neuroplasticity and axonogenesis in the NAc supported CP prevention. Thus, our study shed some light in the gene expression changes in the NAc underlying CP susceptibility, however, our study was not able to investigate deeply those biological processes to suggest and confirm a key role of some specific target gene in the CP susceptibility. Nevertheless, our findings suggest novel insights and potential biologic processes involving the effect of HFD and sedentary behavior in promoting CP susceptibility, as well as, the crucial role of voluntary PA to prevent CP.

## 6. Limitations

One of the limitations of our study was the lack of female mice groups to investigate the effect of HFD and sedentary behavior in promoting CP susceptibility, as well as the effect of voluntary PA to prevent it. Moreover, the lack of additional molecular approaches, such as knockout or knockdown, to test and/or confirm the role of some target genes diminished our odds of establishing causality and the direct effects of variables studied. Despite these limitations, we support the idea that the strict methodological and statistical approaches employed here, as well as findings reported fulfilled crucial scientific questions and provide new avenues for further study.

## Supporting information

Supplementary_R-script

## 7. Acknowledgments

We thank the eScience Institute of the University of Washington and the Hyak HPC team for providing access to the Hyak Supercomputer for RNA-Seq data analysis. We thank Professor Dennys Esper Cyntra and the Laboratory of Nutritional Genomic from the School of Applied Science of the University of Campinas to provide the high-fat diet used in our study. We especially thank the veterinary technician Cesar Eduardo Bissoto for careful attention to the mice in the vivarium.

## 9. Author Contributions

The contributions of authors were organized by: i) conception of the project (AFB, CRS, ASV, CAP); ii) design of the study (AFB, CRS, ASV); iii) acquisition of the data (AFB, IJMB, MPJr., and GGZ); iv) analysis and interpretations of the data (AFB, ASV, and CRS); v) NP and ASV statistical and script programming support for RNA-Seq data analysis; vi) drafting the manuscript (AFB and CRS) and vii) final version approved to be submitted (AFB, IJMB, MPJr., GGZ, CHT, CAP, NP, ASV and CRS). This research was developed with laboratory resources and equipment of the CRS, ASV, CHT, and CAP.

## References

Abarca-Gómez, L., Abdeen, Z. A., Hamid, Z. A., Abu-Rmeileh, N. M., Acosta-Cazares, B., Acuin, C., et al. (2017). Worldwide trends in body-mass index, underweight, overweight, and obesity from 1975 to 2016: a pooled analysis of 2416 population-based measurement studies in 128·9 million children, adolescents, and adults. Lancet 390, 2627–2642. doi:10.1016/S0140-6736(17)32129-3.

American Veterinary Medical Association, A. (2013). AVMA Guidelines for the Euthanasia of Animal: 2013., ed. A. V. M. Association Schaumburg, IL: Elsevier Available at: https://www.avma.org/kb/policies/documents/euthanasia.pdf.

Apkarian, A. V., Baliki, M. N., and Geha, P. Y. (2009). Towards a theory of chronic pain. Prog. Neurobiol. 87, 81–97. doi:10.1016/j.pneurobio.2008.09.018.

Apkarian, A. V., Hashmi, J. A., and Baliki, M. N. (2011). Pain and the brain: specificity and plasticity of the brain in clinical chronic pain. Pain 152, S49–64. doi:10.1016/j.pain.2010.11.010.

Benarroch, E. E. (2016). Involvement of the nucleus accumbens and dopamine system in chronic pain. Neurology 87, 1720–1726. doi:10.1212/WNL.0000000000003243.

Bjursell, M., Gerdin, A.-K., Lelliott, C. J., Egecioglu, E., Elmgren, A., Törnell, J., et al. (2008). Acutely reduced locomotor activity is a major contributor to Western diet-induced obesity in mice. Am. J. Physiol. Metab. 294, E251–E260. doi:10.1152/ajpendo.00401.2007.

Blighe, K. (2019). EnhancedVolcano: Publication-ready volcano plots with enhanced colouring and labeling. doi:10.18129/B9.bioc.EnhancedVolcano.

Breivik, H., Eisenberg, E., and O’Brien, T. (2013). The individual and societal burden of chronic pain in Europe: The case for strategic prioritisation and action to improve knowledge and availability of appropriate care. BMC Public Health 13. doi:10.1186/1471-2458-13-1229.

Castellani, L., Root-Mccaig, J., Frendo-Cumbo, S., Beaudoin, M.-S., and Wright, D. C. (2014). Exercise training protects against an acute inflammatory insult in mouse epididymal adipose tissue. J. Appl. Physiol. 116, 1272–1280. doi:10.1152/japplphysiol.00074.2014.

Chen, W., Wang, H. J., Shang, N. N., Liu, J., Li, J., Tang, D. H., et al. (2017). Moderate intensity treadmill exercise alters food preference via dopaminergic plasticity of ventral tegmental area-nucleus accumbens in obese mice. Neurosci. Lett. 641, 56–61. doi:10.1016/j.neulet.2017.01.055.

Cintra, D. E., Ropelle, E. R., Moraes, J. C., Pauli, J. R., Morari, J., de Souza, C. T., et al. (2012). Unsaturated Fatty Acids Revert Diet-Induced Hypothalamic Inflammation in Obesity. PLoS One 7, e30571. doi:10.1371/journal.pone.0030571.

Cooper, L., Ryan, C. G., Ells, L. J., Hamilton, S., Atkinson, G., Cooper, K., et al. (2018a). Weight loss interventions for adults with overweight/obesity and chronic musculoskeletal pain: a mixed methods systematic review. Obes. Rev. 19, 989–1007. doi:10.1111/obr.12686.

Cooper, M. A., O’Meara, B., Jack, M. M., Elliot, D., Lamb, B., Khan, Z. W., et al. (2018b). Intrinsic Activity of C57BL/6 Substrains Associates with High-Fat Diet-Induced Mechanical Sensitivity in Mice. J. Pain. doi:10.1016/j.jpain.2018.05.005.

de Jong, J. M. A., Larsson, O., Cannon, B., and Nedergaard, J. (2015). A stringent validation of mouse adipose tissue identity markers. Am. J. Physiol. Metab. 308, E1085–E1105. doi:10.1152/ajpendo.00023.2015.

Descalzi, G., Mitsi, V., Purushothaman, I., Gaspari, S., Avrampou, K., Loh, Y.-H. E., et al. (2017). Neuropathic pain promotes adaptive changes in gene expression in brain networks involved in stress and depression. Sci. Signal. 10, eaaj1549. doi:10.1126/scisignal.aaj1549.

Deuis, J. R., Dvorakova, L. S., and Vetter, I. (2017). Methods Used to Evaluate Pain Behaviors in Rodents. Front. Mol. Neurosci. 10, 1–17. doi:10.3389/fnmol.2017.00284.

Dias, E. V., Sartori, C. R., Marião, P. R., Vieira, A. S., Camargo, L. C., Athie, M. C. P., et al. (2015). Nucleus accumbens dopaminergic neurotransmission switches its modulatory action in chronification of inflammatory hyperalgesia. Eur. J. Neurosci. 42, 2380–2389. doi:10.1111/ejn.13015.

Doan, L., Manders, T., and Wang, J. (2015). Neuroplasticity Underlying the Comorbidity of Pain and Depression. Neural Plast. 2015, 1–16. doi:10.1155/2015/504691.

Dobin, A., Davis, C. A., Schlesinger, F., Drenkow, J., Zaleski, C., Jha, S., et al. (2013). STAR: ultrafast universal RNA-seq aligner. Bioinformatics 29, 15–21. doi:10.1093/bioinformatics/bts635.

DosSantos, M. F., Moura, B. de S., and DaSilva, A. F. (2017). Reward circuitry plasticity in pain perception and modulation. Front. Pharmacol. 8, 1–13. doi:10.3389/fphar.2017.00790.

Fernandes, J., Arida, R. M., and Gomez-Pinilla, F. (2017). Physical exercise as an epigenetic modulator of brain plasticity and cognition. Neurosci. Biobehav. Rev. 80, 443–456. doi:10.1016/j.neubiorev.2017.06.012.

Ferreira, S. H., Lorenzetti, B. B., and De Campos, D. I. (1990). Induction, blockade and restoration of a persistent hypersensitive state. Pain 42, 365–71. Available at: http://www.ncbi.nlm.nih.gov/pubmed/2174528.

Franklin, K. B., and Paxinos, G. (1996). The Mouse Brain in Stereotaxic Coordinates. San Diego: Academic Press, Inc.

Friend, D. M., Devarakonda, K., O’Neal, T. J., Skirzewski, M., Papazoglou, I., Kaplan, A. R., et al. (2017). Basal Ganglia Dysfunction Contributes to Physical Inactivity in Obesity. Cell Metab. 25, 312–321. doi:10.1016/j.cmet.2016.12.001.

Funkat, A., Massa, C. M., Jovanovska, V., Proietto, J., and Andrikopoulos, S. (2004). Metabolic Adaptations of Three Inbred Strains of Mice (C57BL/6, DBA/2, and 129T2) in Response to a High-Fat Diet. J. Nutr. 134, 3264–3269. doi:10.1093/jn/134.12.3264.

Gaskin, D. J., and Richard, P. (2012). The Economic Costs of Pain in the United States. J. Pain 13, 715–724. doi:10.1016/j.jpain.2012.03.009.

Geneen, L. J., Moore, R. A., Clarke, C., Martin, D., Colvin, L. A., and Smith, B. H. (2017). “Physical activity and exercise for chronic pain in adults: an overview of Cochrane Reviews,” in Cochrane Database of Systematic Reviews, ed. L. J. Geneen (Chichester, UK: John Wiley & Sons, Ltd), CD011279. doi:10.1002/14651858.CD011279.pub2.

Hu, S., Wang, L., Yang, D., Li, L., Togo, J., Wu, Y., et al. (2018a). Dietary Fat, but Not Protein or Carbohydrate, Regulates Energy Intake and Causes Adiposity in Mice. Cell Metab., 415–431. doi:10.1016/j.cmet.2018.06.010.

Hu, Z. J., Han, W., Cao, C. Q., Mao-Ying, Q. L., Mi, W. L., and Wang, Y. Q. (2018b). Peripheral Leptin Signaling Mediates Formalin-Induced Nociception. Neurosci. Bull. 34, 321–329. doi:10.1007/s12264-017-0194-2.

Jayaraman, A., Lent-Schochet, D., and Pike, C. J. (2014). Diet-induced obesity and low testosterone increase neuroinflammation and impair neural function. J. Neuroinflammation 11, 162. doi:10.1186/s12974-014-0162-y.

Jeong, H., Moye, L. S., Southey, B. R., Hernandez, A. G., Dripps, I., Romanova, E. V., et al. (2018). Gene Network Dysregulation in the Trigeminal Ganglia and Nucleus Accumbens of a Model of Chronic Migraine-Associated Hyperalgesia. Front. Syst. Neurosci. 12, 1–19. doi:10.3389/fnsys.2018.00063.

Kleinert, M., Clemmensen, C., Hofmann, S. M., Moore, M. C., Renner, S., Woods, S. C., et al. (2018). Animal models of obesity and diabetes mellitus. Nat. Rev. Endocrinol. 14, 140–162. doi:10.1038/nrendo.2017.161.

Koyanagi, A., Stubbs, B., and Vancampfort, D. (2018). Correlates of sedentary behavior in the general population: A cross-sectional study using nationally representative data from six low- and middle-income countries. PLoS One 13, e0202222. doi:10.1371/journal.pone.0202222.

Kuner, R., and Flor, H. (2016). Structural plasticity and reorganisation in chronic pain. Nat. Rev. Neurosci. 18, 20–30. doi:10.1038/nrn.2016.162.

Leung, A., Gregory, N. S., Allen, L. A. H., and Sluka, K. A. (2016). Regular physical activity prevents chronic pain by altering resident muscle macrophage phenotype and increasing interleukin-10 in mice. Pain 157, 70–79. doi:10.1097/j.pain.0000000000000312.

Li, X., Kang, L., Li, G., Zeng, H., Zhang, L., Ling, X., et al. (2013). Intrathecal leptin inhibits expression of the P2X2/3 receptors and alleviates neuropathic pain induced by chronic constriction sciatic nerve injury. Mol. Pain 9, 65. doi:10.1186/1744-8069-9-65.

Liang, Y. J., Feng, S. Y., Qi, Y. P., Li, K., Jin, Z. R., Jing, H. B., et al. (2019). Contribution of microglial reaction to increased nociceptive responses in high-fat-diet (HFD)-induced obesity in male mice. Brain. Behav. Immun. 80, 777–792. doi:10.1016/j.bbi.2019.05.026.

Lima, L. V., Abner, T. S. S., and Sluka, K. A. (2017). Does exercise increase or decrease pain? Central mechanisms underlying these two phenomena. J. Physiol. 00, 1–10. doi:10.1113/JP273355.

Lourenco, M. V, Frozza, R. L., de Freitas, G. B., Zhang, H., Kincheski, G. C., Ribeiro, F. C., et al. (2019). Exercise-linked FNDC5/irisin rescues synaptic plasticity and memory defects in Alzheimer’s models. Nat. Med. 25, 165–175. doi:10.1038/s41591-018-0275-4.

Love, M. I., Huber, W., and Anders, S. (2014). Moderated estimation of fold change and dispersion for RNA-seq data with DESeq2. Genome Biol. 15, 1–21. doi:10.1186/s13059-014-0550-8.

Mansour, A. R., Farmer, M. A., Baliki, M. N., and Apkarian, A. V. (2014). Chronic pain: The role of learning and brain plasticity. Restor. Neurol. Neurosci. 32, 129–139. doi:10.3233/RNN-139003.

Martikainen, I. K., Nuechterlein, E. B., Pecina, M., Love, T. M., Cummiford, C. M., Green, C. R., et al. (2015). Chronic Back Pain Is Associated with Alterations in Dopamine Neurotransmission in the Ventral Striatum. J. Neurosci. 35, 9957–9965. doi:10.1523/JNEUROSCI.4605-14.2015.

Martinov, T., Mack, M., Sykes, A., and Chatterjea, D. (2013). Measuring changes in tactile sensitivity in the hind paw of mice using an electronic von Frey apparatus. J. Vis. Exp., e51212. doi:10.3791/51212.

Mayer, S., Spickschen, J., Stein, K. V., Crevenna, R., Dorner, T. E., and Simon, J. (2019). The societal costs of chronic pain and its determinants: The case of Austria. PLoS One 14, 1–18. doi:10.1371/journal.pone.0213889.

Montgomery, M. K., Hallahan, N. L., Brown, S. H., Liu, M., Mitchell, T. W., Cooney, G. J., et al. (2013). Mouse strain-dependent variation in obesity and glucose homeostasis in response to high-fat feeding. Diabetologia 56, 1129–1139. doi:10.1007/s00125-013-2846-8.

Paley, C. A., and Johnson, M. I. (2016). Physical Activity to Reduce Systemic Inflammation Associated With Chronic Pain and Obesity. Clin. J. Pain 32, 365–370. doi:10.1097/AJP.0000000000000258.

Ren, W., Centeno, M. V., Berger, S., Wu, Y., Na, X., Liu, X., et al. (2015). The indirect pathway of the nucleus accumbens shell amplifies neuropathic pain. Nat. Neurosci. 19, 220–222. doi:10.1038/nn.4199.

Rocha-Rodrigues, S., Gonçalves, I. O., Beleza, J., Ascensão, A., and Magalhães, J. (2018). Physical exercise mitigates high-fat diet-induced adiposopathy and related endocrine alterations in an animal model of obesity. J. Physiol. Biochem. 74, 235–246. doi:10.1007/s13105-018-0609-1.

Rossi, H. L., Luu, A. K. S., Devilbiss, J. L., and Recober, A. (2013). Obesity increases nociceptive activation of the trigeminal system. Eur. J. Pain (United Kingdom) 17, 649–653. doi:10.1002/j.1532-2149.2012.00230.x.

Sachs, D., Villarreal, C., Cunha, F., Parada, C., and Ferreira, S. (2009). The role of PKA and PKCε pathways in prostaglandin E2-mediated hypernociception. Br. J. Pharmacol. 156, 826–834. doi:10.1111/j.1476-5381.2008.00093.x.

Salamone, J. D., and Correa, M. (2012). The Mysterious Motivational Functions of Mesolimbic Dopamine. Neuron 76, 470–485. doi:10.1016/j.neuron.2012.10.021.

Salamone, J. D., Pardo, M., Yohn, S. E., López-Cruz, L., SanMiguel, N., and Correa, M. (2015). “Mesolimbic Dopamine and the Regulation of Motivated Behavior,” in Current Topics in Behavioral Neurosciences, 231–257. doi:10.1007/7854_2015_383.

Schwartz, N., Miller, C., and Fields, H. L. (2017). Cortico-Accumbens Regulation of Approach-Avoidance Behavior Is Modified by Experience and Chronic Pain. Cell Rep. 19, 1522–1531. doi:10.1016/j.celrep.2017.04.073.

Sluka, K. A., O’Donnell, J. M., Danielson, J., and Rasmussen, L. A. (2013). Regular physical activity prevents development of chronic pain and activation of central neurons. J. Appl. Physiol. 114, 725–733. doi:10.1152/japplphysiol.01317.2012.

Song, Z., Xie, W., Chen, S., Strong, J. A., Print, M. S., Wang, J. I., et al. (2017). High-fat diet increases pain behaviors in rats with or without obesity. Sci. Rep. 7, 10350. doi:10.1038/s41598-017-10458-z.

Souza, G. R., Cunha, T. M., Silva, R. L., Lotufo, C. M., Verri, W. A., Funez, M. I., et al. (2015). Involvement of nuclear factor kappa B in the maintenance of persistent inflammatory hypernociception. Pharmacol. Biochem. Behav. 134, 49–56. doi:10.1016/j.pbb.2015.04.005.

Starobova, H., Himaya, S. W. A., Lewis, R. J., and Vetter, I. (2018). Transcriptomics in pain research: insights from new and old technologies. Mol. Omi. 14, 389–404. doi:10.1039/c8mo00181b.

Torrance, N., Smith, B. H., Bennett, M. I., and Lee, A. J. (2006). The Epidemiology of Chronic Pain of Predominantly Neuropathic Origin. Results From a General Population Survey. J. Pain 7, 281–289. doi:10.1016/j.jpain.2005.11.008.

Van Hecke, O., Austin, S. K., Khan, R. A., Smith, B. H., and Torrance, N. (2014). Neuropathic pain in the general population: A systematic review of epidemiological studies. Pain 155, 654–662. doi:10.1016/j.pain.2013.11.013.

Van Hecke, O., Torrance, N., and Smith, B. H. (2013). Chronic pain epidemiology and its clinical relevance. Br. J. Anaesth. 111, 13–18. doi:10.1093/bja/aet123.

Vellers, H. L., Letsinger, A. C., Walker, N. R., Granados, J. Z., and Lightfoot, J. T. (2017). High fat high sugar diet reduces voluntary wheel running in mice independent of sex hormone involvement. Front. Physiol. 8, 1–14. doi:10.3389/fphys.2017.00628.

Villarreal, C. F., Funez, M. I., Figueiredo, F., Cunha, F. Q., Parada, C. A., and Ferreira, S. H. (2009). Acute and persistent nociceptive paw sensitisation in mice: The involvement of distinct signalling pathways. Life Sci. 85, 822–829. doi:10.1016/j.lfs.2009.10.018.

Williams, L. M., Campbell, F. M., Drew, J. E., Koch, C., Hoggard, N., Rees, W. D., et al. (2014). The Development of Diet-Induced Obesity and Glucose Intolerance in C57Bl/6 Mice on a High-Fat Diet Consists of Distinct Phases. PLoS One 9, e106159. doi:10.1371/journal.pone.0106159.

Woolf, C. J. (2000). Neuronal Plasticity: Increasing the Gain in Pain. Science (80-.). 288, 1765–1768. doi:10.1126/science.288.5472.1765.

Yang, Y., Smith, D. L., Keating, K. D., Allison, D. B., and Nagy, T. R. (2014). Variations in body weight, Food Intake and body composition after long-term high-fat diet feeding in C57BL/6J mice. Obesity 22, 2147–2155. doi:10.1002/oby.20811.

Zhang, P., Moye, L. S., Southey, B. R., Dripps, I., Sweedler, J. V., Pradhan, A., et al. (2019). Opioid-Induced Hyperalgesia Is Associated with Dysregulation of Circadian Rhythm and Adaptive Immune Pathways in the Mouse Trigeminal Ganglia and Nucleus Accumbens. Mol. Neurobiol. doi:10.1007/s12035-019-01650-5.

