## Supplementary_R-script for "Physical Activity Induces Nucleus Accumbens Genes Expression Changes Preventing Chronic Pain Susceptibility Promoted by High-Fat Diet and Sedentary Behavior in Mice"

```
#References:
# 1) Love MI, Anders S, Kim V, Huber W (2016). RNA-Seq workflow: gene-level exploratory analysis and
differential
# expression. F1000Research 4:1070 Available at: https://f1000research.com/articles/4-1070/v2.

# 2) Love MI, Huber W, Anders S (2014). Moderated estimation of fold change and dispersion for RNA-seq data
with
# DESeq2. Genome Biol 15:1–21

#(https://www.bioconductor.org/packages/devel/bioc/vignettes/EnhancedVolcano/inst/doc/EnhancedVolcano.html)

################ Start ##############

library(DESeq2)
library(ggplot2)
library(pheatmap)
library(RColorBrewer)
library(genefilter)
library(EnhancedVolcano)
library(gridExtra)
library(gridBase)
library(STRINGdb)


file <- read.csv("/home/afb/AL3_data/AL3_data.csv", sep = ",", row.names = "gene")
head (file)

AL3_matrix <- as.matrix(file)#cria uma matrix para ser lida pelo DESeq
head(AL3_matrix)
coldata_AL3 <- read.csv("/home/afb/AL3_data/col_data_AL3.csv", sep = ",", row.names = "samples")

########
coldata_AL3 <- coldata_AL3[,c("treat", "rw", "diet")]
########

head (coldata_AL3)
summary (coldata_AL3)
all(rownames(coldata_AL3) %in% colnames(AL3_matrix))
all(rownames(coldata_AL3) == colnames(AL3_matrix))

############################
# 1.3.3 Count matrix input

dds <- DESeqDataSetFromMatrix(countData = AL3_matrix, colData = coldata_AL3, design = ~ treat + rw + diet)
dds

# 1.3.6 Pre-filtering and rlog normalization
dds <- dds[ rowSums(counts(dds)) > 0.9, ]
dds
colData(dds)
vsd_dds <- vst(dds)
#############################################


######### DESeq2 #####################################
#1.4 Differential expression analysis
dds <- DESeq(dds)
#######################################

#----------------------------------------#--------------------------------------------------#--------------------------------------------------#
colData(dds)
ddsIF_3 <- dds
```

```
ddsIF_3$group <- factor(paste0(ddsIF_3$rw, ddsIF_3$diet, ddsIF_3$treat))
design(ddsIF_3) <- ~ group
ddsIF_3 <- DESeq(ddsIF_3)
resultsNames(ddsIF_3)

#---- write table---#
write.csv(as.data.frame(raw_ddsIF_3), file = "/home/afb/AL3_data/raw_ddsIF_3.csv")
write.csv(as.data.frame(means_ddsIF_3), file = "/home/afb/AL3_data/means_ddsIF_3.csv")
write.csv(as.data.frame(cooks_ddsIF_3), file = "/home/afb/AL3_data/cooks_ddsIF_3.csv")
#---------------------#

#### 1)  SED.SD.SAL vs PA.HFD.SAL ######
SED.SD.SAL_PA.HFD.SAL <- results(ddsIF_3, contrast = c("group", "SEDSDSAL", "PAHFDSAL" ))
SED.SD.SAL_PA.HFD.SAL <- SED.SD.SAL_PA.HFD.SAL[order(SED.SD.SAL_PA.HFD.SAL$padj),]
SED.SD.SAL_PA.HFD.SAL
summary(SED.SD.SAL_PA.HFD.SAL)
sum(SED.SD.SAL_PA.HFD.SAL$padj < 0.1, na.rm = TRUE)
sum(SED.SD.SAL_PA.HFD.SAL$padj < 0.05, na.rm = TRUE)
write.csv(as.data.frame(SED.SD.SAL_PA.HFD.SAL), file =
"/home/afb/AL3_data/AL3_SED.SD.SAL_PA.HFD.SAL.csv")

#### 2)  SED.SD.SAL vs PA.HFD.PGE ######
SED.SD.SAL_PA.HFD.PGE <- results(ddsIF_3, contrast = c("group", "SEDSDSAL", "PAHFDPGE" ))
SED.SD.SAL_PA.HFD.PGE <- SED.SD.SAL_PA.HFD.PGE[order(SED.SD.SAL_PA.HFD.PGE$padj),]
SED.SD.SAL_PA.HFD.PGE
summary(SED.SD.SAL_PA.HFD.PGE)
sum(SED.SD.SAL_PA.HFD.PGE$padj < 0.1, na.rm = TRUE)
sum(SED.SD.SAL_PA.HFD.PGE$padj < 0.05, na.rm = TRUE)
write.csv(as.data.frame(SED.SD.SAL_PA.HFD.PGE), file =
"/home/afb/AL3_data/AL3_SED.SD.SAL_PA.HFD.PGE.csv")

#### 3)  SED.SD.PGE vs PA.HFD.SAL ######
SED.SD.PGE_PA.HFD.SAL <- results(ddsIF_3, contrast = c("group", "SEDSDPGE", "PAHFDSAL" ))
SED.SD.PGE_PA.HFD.SAL <- SED.SD.PGE_PA.HFD.SAL[order(SED.SD.PGE_PA.HFD.SAL$padj),]
SED.SD.PGE_PA.HFD.SAL
summary(SED.SD.PGE_PA.HFD.SAL)
sum(SED.SD.PGE_PA.HFD.SAL$padj < 0.1, na.rm = TRUE)
sum(SED.SD.PGE_PA.HFD.SAL$padj < 0.05, na.rm = TRUE)
write.csv(as.data.frame(SED.SD.PGE_PA.HFD.SAL), file =
"/home/afb/AL3_data/AL3_SED.SD.PGE_PA.HFD.SAL.csv")

#### 4)  SED.SD.PGE vs PA.HFD.PGE ######
SED.SD.PGE_PA.HFD.PGE <- results(ddsIF_3, contrast = c("group", "SEDSDPGE", "PAHFDPGE" ))
SED.SD.PGE_PA.HFD.PGE <- SED.SD.PGE_PA.HFD.PGE[order(SED.SD.PGE_PA.HFD.PGE$padj),]
SED.SD.PGE_PA.HFD.PGE
summary(SED.SD.PGE_PA.HFD.PGE)
sum(SED.SD.PGE_PA.HFD.PGE$padj < 0.1, na.rm = TRUE)
sum(SED.SD.PGE_PA.HFD.PGE$padj < 0.05, na.rm = TRUE)
write.csv(as.data.frame(SED.SD.PGE_PA.HFD.PGE), file =
"/home/afb/AL3_data/AL3_SED.SD.PGE_PA.HFD.PGE.csv")

#### 5)  SED.SD.SAL vs PA.SD.SAL ######
SED.SD.SAL_PA.SD.SAL <- results(ddsIF_3, contrast = c("group", "SEDSDSAL", "PASDSAL" ))
SED.SD.SAL_PA.SD.SAL <- SED.SD.SAL_PA.SD.SAL[order(SED.SD.SAL_PA.SD.SAL$padj),]
SED.SD.SAL_PA.SD.SAL
summary(SED.SD.SAL_PA.SD.SAL)
sum(SED.SD.SAL_PA.SD.SAL$padj < 0.1, na.rm = TRUE)
sum(SED.SD.SAL_PA.SD.SAL$padj < 0.05, na.rm = TRUE)
write.csv(as.data.frame(SED.SD.SAL_PA.SD.SAL), file =
"/home/afb/AL3_data/AL3_SED.SD.SAL_PA.SD.SAL.csv")

#### 6)  SED.SD.SAL vs PA.SD.PGE ######
```

```
SED.SD.SAL_PA.SD.PGE <- results(ddsIF_3, contrast = c("group", "SEDSDSAL", "PASDPGE" ))
SED.SD.SAL_PA.SD.PGE <- SED.SD.SAL_PA.SD.PGE[order(SED.SD.SAL_PA.SD.PGE$padj),]
SED.SD.SAL_PA.SD.PGE
summary(SED.SD.SAL_PA.SD.PGE)
sum(SED.SD.SAL_PA.SD.PGE$padj < 0.1, na.rm = TRUE)
sum(SED.SD.SAL_PA.SD.PGE$padj < 0.05, na.rm = TRUE)
write.csv(as.data.frame(SED.SD.SAL_PA.SD.PGE), file =
"/home/afb/AL3_data/AL3_SED.SD.SAL_PA.SD.PGE.csv")

#### 7)  SED.SD.PGE vs PA.SD.SAL ######
SED.SD.PGE_PA.SD.SAL <- results(ddsIF_3, contrast = c("group", "SEDSDPGE", "PASDSAL" ))
SED.SD.PGE_PA.SD.SAL <- SED.SD.PGE_PA.SD.SAL[order(SED.SD.PGE_PA.SD.SAL$padj),]
SED.SD.PGE_PA.SD.SAL
summary(SED.SD.PGE_PA.SD.SAL)
sum(SED.SD.PGE_PA.SD.SAL$padj < 0.1, na.rm = TRUE)
sum(SED.SD.PGE_PA.SD.SAL$padj < 0.05, na.rm = TRUE)
write.csv(as.data.frame(SED.SD.PGE_PA.SD.SAL), file =
"/home/afb/AL3_data/AL3_SED.SD.PGE_PA.SD.SAL.csv")

#### 8)  SED.SD.PGE vs PA.SD.PGE ######
SED.SD.PGE_PA.SD.PGE <- results(ddsIF_3, contrast = c("group", "SEDSDPGE", "PASDPGE" ))
SED.SD.PGE_PA.SD.PGE <- SED.SD.PGE_PA.SD.PGE[order(SED.SD.PGE_PA.SD.PGE$padj),]
SED.SD.PGE_PA.SD.PGE
summary(SED.SD.PGE_PA.SD.PGE)
sum(SED.SD.PGE_PA.SD.PGE$padj < 0.1, na.rm = TRUE)
sum(SED.SD.PGE_PA.SD.PGE$padj < 0.05, na.rm = TRUE)
write.csv(as.data.frame(SED.SD.PGE_PA.SD.PGE), file =
"/home/afb/AL3_data/AL3_SED.SD.PGE_PA.SD.PGE.csv")

#### 9)  SED.HFD.SAL vs PA.SD.SAL ######
SED.HFD.SAL_PA.SD.SAL <- results(ddsIF_3, contrast = c("group", "SEDHFDSAL", "PASDSAL" ))
SED.HFD.SAL_PA.SD.SAL <- SED.HFD.SAL_PA.SD.SAL[order(SED.HFD.SAL_PA.SD.SAL$padj),]
SED.HFD.SAL_PA.SD.SAL
summary(SED.HFD.SAL_PA.SD.SAL)
sum(SED.HFD.SAL_PA.SD.SAL$padj < 0.1, na.rm = TRUE)
sum(SED.HFD.SAL_PA.SD.SAL$padj < 0.05, na.rm = TRUE)
write.csv(as.data.frame(SED.HFD.SAL_PA.SD.SAL), file =
"/home/afb/AL3_data/AL3_SED.HFD.SAL_PA.SD.SAL.csv")

#### 10)  SED.HFD.SAL vs PA.SD.PGE ######
SED.HFD.SAL_PA.SD.PGE <- results(ddsIF_3, contrast = c("group", "SEDHFDSAL", "PASDPGE" ))
SED.HFD.SAL_PA.SD.PGE <- SED.HFD.SAL_PA.SD.PGE[order(SED.HFD.SAL_PA.SD.PGE$padj),]
SED.HFD.SAL_PA.SD.PGE
summary(SED.HFD.SAL_PA.SD.PGE)
sum(SED.HFD.SAL_PA.SD.PGE$padj < 0.1, na.rm = TRUE)
sum(SED.HFD.SAL_PA.SD.PGE$padj < 0.05, na.rm = TRUE)
write.csv(as.data.frame(SED.HFD.SAL_PA.SD.PGE), file =
"/home/afb/AL3_data/AL3_SED.HFD.SAL_PA.SD.PGE.csv")

#### 11)  SED.HFD.PGE vs PA.SD.SAL ######
SED.HFD.PGE_PA.SD.SAL <- results(ddsIF_3, contrast = c("group", "SEDHFDPGE", "PASDSAL" ))
SED.HFD.PGE_PA.SD.SAL <- SED.HFD.PGE_PA.SD.SAL[order(SED.HFD.PGE_PA.SD.SAL$padj),]
SED.HFD.PGE_PA.SD.SAL
summary(SED.HFD.PGE_PA.SD.SAL)
sum(SED.HFD.PGE_PA.SD.SAL$padj < 0.1, na.rm = TRUE)
sum(SED.HFD.PGE_PA.SD.SAL$padj < 0.05, na.rm = TRUE)
write.csv(as.data.frame(SED.HFD.PGE_PA.SD.SAL), file =
"/home/afb/AL3_data/AL3_SED.HFD.PGE_PA.SD.SAL.csv")

#### 12)  SED.HFD.PGE vs PA.SD.PGE ######
SED.HFD.PGE_PA.SD.PGE <- results(ddsIF_3, contrast = c("group", "SEDHFDPGE", "PASDPGE" ))
SED.HFD.PGE_PA.SD.PGE <- SED.HFD.PGE_PA.SD.PGE[order(SED.HFD.PGE_PA.SD.PGE$padj),]
```

```
SED.HFD.PGE_PA.SD.PGE
summary(SED.HFD.PGE_PA.SD.PGE)
sum(SED.HFD.PGE_PA.SD.PGE$padj < 0.1, na.rm = TRUE)
sum(SED.HFD.PGE_PA.SD.PGE$padj < 0.05, na.rm = TRUE)
write.csv(as.data.frame(SED.HFD.PGE_PA.SD.PGE), file =
"/home/afb/AL3_data/AL3_SED.HFD.PGE_PA.SD.PGE.csv")

#### 13)  SED.SD.SAL vs SED.SD.PGE ######
SED.SD.SAL_SED.SD.PGE <- results(ddsIF_3, contrast = c("group", "SEDSDSAL", "SEDSDPGE" ))
SED.SD.SAL_SED.SD.PGE <- SED.SD.SAL_SED.SD.PGE[order(SED.SD.SAL_SED.SD.PGE$padj),]
SED.SD.SAL_SED.SD.PGE
summary(SED.SD.SAL_SED.SD.PGE)
sum(SED.SD.SAL_SED.SD.PGE$padj < 0.1, na.rm = TRUE)
sum(SED.SD.SAL_SED.SD.PGE$padj < 0.05, na.rm = TRUE)
write.csv(as.data.frame(SED.SD.SAL_SED.SD.PGE), file =
"/home/afb/AL3_data/AL3_SED.SD.SAL_SED.SD.PGE.csv")

#### 14)  SED.HFD.SAL vs SED.HFD.PGE ######
SED.HFD.SAL_SED.HFD.PGE <- results(ddsIF_3, contrast = c("group", "SEDHFDSAL", "SEDHFDPGE" ))
SED.HFD.SAL_SED.HFD.PGE <-
SED.HFD.SAL_SED.HFD.PGE[order(SED.HFD.SAL_SED.HFD.PGE$padj),]
SED.HFD.SAL_SED.HFD.PGE
summary(SED.HFD.SAL_SED.HFD.PGE)
sum(SED.HFD.SAL_SED.HFD.PGE$padj < 0.1, na.rm = TRUE)
sum(SED.HFD.SAL_SED.HFD.PGE$padj < 0.05, na.rm = TRUE)
write.csv(as.data.frame(SED.HFD.SAL_SED.HFD.PGE), file =
"/home/afb/AL3_data/AL3_SED.HFD.SAL_SED.HFD.PGE.csv")

#### 15)  SED.SD.SAL vs SED.HFD.SAL ######
SED.SD.SAL_SED.HFD.SAL <- results(ddsIF_3, contrast = c("group", "SEDSDSAL", "SEDHFDSAL" ))
SED.SD.SAL_SED.HFD.SAL <- SED.SD.SAL_SED.HFD.SAL[order(SED.SD.SAL_SED.HFD.SAL$padj),]
SED.SD.SAL_SED.HFD.SAL
summary(SED.SD.SAL_SED.HFD.SAL)
sum(SED.SD.SAL_SED.HFD.SAL$padj < 0.1, na.rm = TRUE)
sum(SED.SD.SAL_SED.HFD.SAL$padj < 0.05, na.rm = TRUE)
write.csv(as.data.frame(SED.SD.SAL_SED.HFD.SAL), file =
"/home/afb/AL3_data/AL3_SED.SD.SAL_SED.HFD.SAL.csv")

#### 16)  SED.SD.PGE vs SED.HFD.PGE ######
SED.SD.PGE_SED.HFD.PGE <- results(ddsIF_3, contrast = c("group", "SEDSDPGE", "SEDHFDPGE" ))
SED.SD.PGE_SED.HFD.PGE <- SED.SD.PGE_SED.HFD.PGE[order(SED.SD.PGE_SED.HFD.PGE$padj),]
SED.SD.PGE_SED.HFD.PGE
summary(SED.SD.PGE_SED.HFD.PGE)
sum(SED.SD.PGE_SED.HFD.PGE$padj < 0.1, na.rm = TRUE)
sum(SED.SD.PGE_SED.HFD.PGE$padj < 0.05, na.rm = TRUE)
write.csv(as.data.frame(SED.SD.PGE_SED.HFD.PGE), file =
"/home/afb/AL3_data/AL3_SED.SD.PGE_SED.HFD.PGE.csv")

#### 17)  PA.SD.SAL vs PA.SD.PGE ######
PA.SD.SAL_PA.SD.PGE <- results(ddsIF_3, contrast = c("group", "PASDSAL", "PASDPGE" ))
PA.SD.SAL_PA.SD.PGE <- PA.SD.SAL_PA.SD.PGE[order(PA.SD.SAL_PA.SD.PGE$padj),]
PA.SD.SAL_PA.SD.PGE
summary(PA.SD.SAL_PA.SD.PGE)
sum(PA.SD.SAL_PA.SD.PGE$padj < 0.1, na.rm = TRUE)
sum(PA.SD.SAL_PA.SD.PGE$padj < 0.05, na.rm = TRUE)
write.csv(as.data.frame(PA.SD.SAL_PA.SD.PGE), file =
"/home/afb/AL3_data/AL3_PA.SD.SAL_PA.SD.PGE.csv")

#### 18)  PA.HFD.SAL vs PA.HFD.PGE ######
PA.HFD.SAL_PA.HFD.PGE <- results(ddsIF_3, contrast = c("group", "PAHFDSAL", "PAHFDPGE" ))
PA.HFD.SAL_PA.HFD.PGE <- PA.HFD.SAL_PA.HFD.PGE[order(PA.HFD.SAL_PA.HFD.PGE$padj),]
PA.HFD.SAL_PA.HFD.PGE
```

```
summary(PA.HFD.SAL_PA.HFD.PGE)
sum(PA.HFD.SAL_PA.HFD.PGE$padj < 0.1, na.rm = TRUE)
sum(PA.HFD.SAL_PA.HFD.PGE$padj < 0.05, na.rm = TRUE)
write.csv(as.data.frame(PA.HFD.SAL_PA.HFD.PGE), file =
"/home/afb/AL3_data/AL3_PA.HFD.SAL_PA.HFD.PGE.csv")

#### 19)  PA.SD.SAL vs PA.HFD.SAL ######
PA.SD.SAL_PA.HFD.SAL <- results(ddsIF_3, contrast = c("group", "PASDSAL", "PAHFDSAL" ))
PA.SD.SAL_PA.HFD.SAL <- PA.SD.SAL_PA.HFD.SAL[order(PA.SD.SAL_PA.HFD.SAL$padj),]
PA.SD.SAL_PA.HFD.SAL
summary(PA.SD.SAL_PA.HFD.SAL)
sum(PA.SD.SAL_PA.HFD.SAL$padj < 0.1, na.rm = TRUE)
sum(PA.SD.SAL_PA.HFD.SAL$padj < 0.05, na.rm = TRUE)
write.csv(as.data.frame(PA.SD.SAL_PA.HFD.SAL), file =
"/home/afb/AL3_data/AL3_PA.SD.SAL_PA.HFD.SAL.csv")

#### 20)  PA.SD.PGE vs PA.HFD.PGE ######
PA.SD.PGE_PA.HFD.PGE <- results(ddsIF_3, contrast = c("group", "PASDPGE", "PAHFDPGE" ))
PA.SD.PGE_PA.HFD.PGE <- PA.SD.PGE_PA.HFD.PGE[order(PA.SD.PGE_PA.HFD.PGE$padj),]
PA.SD.PGE_PA.HFD.PGE
summary(PA.SD.PGE_PA.HFD.PGE)
sum(PA.SD.PGE_PA.HFD.PGE$padj < 0.1, na.rm = TRUE)
sum(PA.SD.PGE_PA.HFD.PGE$padj < 0.05, na.rm = TRUE)
write.csv(as.data.frame(PA.SD.PGE_PA.HFD.PGE), file =
"/home/afb/AL3_data/AL3_PA.SD.PGE_PA.HFD.PGE.csv")

#### 21)  SED.HFD.SAL vs PA.HFD.SAL ######
SED.HFD.SAL_PA.HFD.SAL <- results(ddsIF_3, contrast = c("group", "SEDHFDSAL", "PAHFDSAL" ))
SED.HFD.SAL_PA.HFD.SAL <- SED.HFD.SAL_PA.HFD.SAL[order(SED.HFD.SAL_PA.HFD.SAL$padj),]
SED.HFD.SAL_PA.HFD.SAL
summary(SED.HFD.SAL_PA.HFD.SAL)
sum(SED.HFD.SAL_PA.HFD.SAL$padj < 0.1, na.rm = TRUE)
sum(SED.HFD.SAL_PA.HFD.SAL$padj < 0.05, na.rm = TRUE)
write.csv(as.data.frame(SED.HFD.SAL_PA.HFD.SAL), file =
"/home/afb/AL3_data/AL3_SED.HFD.SAL_PA.HFD.SAL.csv")

#### 22)  SED.HFD.PGE vs PA.HFD.PGE ######
SED.HFD.PGE_PA.HFD.PGE <- results(ddsIF_3, contrast = c("group", "SEDHFDPGE", "PAHFDPGE" ))
SED.HFD.PGE_PA.HFD.PGE <- SED.HFD.PGE_PA.HFD.PGE[order(SED.HFD.PGE_PA.HFD.PGE$padj),]
SED.HFD.PGE_PA.HFD.PGE
summary(SED.HFD.PGE_PA.HFD.PGE)
sum(SED.HFD.PGE_PA.HFD.PGE$padj < 0.1, na.rm = TRUE)
sum(SED.HFD.PGE_PA.HFD.PGE$padj < 0.05, na.rm = TRUE)
mcols(ddsIF_3, use.names = TRUE)[1:4,1:4] # acesso aos valores de dispersão, coeficientes e SE
write.csv(as.data.frame(SED.HFD.PGE_PA.HFD.PGE), file =
"/home/afb/AL3_data/AL3_SED.HFD.PGE_PA.HFD.PGE.csv")

#--------------# Volcano Plot #---------------------------#

EnhancedVolcano(SED.HFD.PGE_PA.HFD.PGE,
        lab = rownames(SED.HFD.PGE_PA.HFD.PGE),
        x = 'log2FoldChange',
        y = 'padj',
        xlim = c(-3, 4),
        ylim = c(0, 3),
        selectLab = c('FALSE'),
        title = 'SED.HFD.PGE_vs_PA.HFD.PGE',
        xlab = bquote(~Log[2]~ 'fold-change'),
        ylab = bquote(~Adjusted~p-value),
        cutoffLineType = 'twodash',
        cutoffLineWidth = 0.6,
        pCutoff = 0.05,
```

```
        FCcutoff = 1.0,
        transcriptPointSize = 2.5,
        transcriptLabSize = 3,
        boxedlabels = FALSE,
        col=c('grey0', 'green3', 'blue1', 'red2'),
        colAlpha = 0.3,
        shape = 20,
        legend=c('Non-significant','Log2 fold-change','Adjust p-value','Adjust p-value & Log2 fold-change'),
        legendPosition = 'right',
        legendLabSize = 12,
        legendIconSize = 5.0,
        drawConnectors = FALSE,
        widthConnectors = 1.0,
        colConnectors = 'black',
        gridlines.major = FALSE,
        gridlines.minor = FALSE)

#--------------# Volcano Plot #---------------------------#

#### 23)  SED.HFD.SAL vs PA.HFD.PGE ######
SED.HFD.SAL_PA.HFD.PGE <- results(ddsIF_3, contrast = c("group", "SEDHFDSAL", "PAHFDPGE" ))
SED.HFD.SAL_PA.HFD.PGE <- SED.HFD.SAL_PA.HFD.PGE[order(SED.HFD.SAL_PA.HFD.PGE$padj),]
SED.HFD.SAL_PA.HFD.PGE
summary(SED.HFD.SAL_PA.HFD.PGE)
sum(SED.HFD.SAL_PA.HFD.PGE$padj < 0.1, na.rm = TRUE)
sum(SED.HFD.SAL_PA.HFD.PGE$padj < 0.05, na.rm = TRUE)
write.csv(as.data.frame(SED.HFD.SAL_PA.HFD.PGE), file =
"/home/afb/AL3_data/AL3_SED.HFD.SAL_PA.HFD.PGE.csv")

#### 24)  SED.HFD.PGE vs PA.HFD.SAL ######
SED.HFD.PGE_PA.HFD.SAL <- results(ddsIF_3, contrast = c("group", "SEDHFDPGE", "PAHFDSAL" ))
SED.HFD.PGE_PA.HFD.SAL <- SED.HFD.PGE_PA.HFD.SAL[order(SED.HFD.PGE_PA.HFD.SAL$padj),]
SED.HFD.PGE_PA.HFD.SAL
summary(SED.HFD.PGE_PA.HFD.SAL)
sum(SED.HFD.PGE_PA.HFD.SAL$padj < 0.1, na.rm = TRUE)
sum(SED.HFD.PGE_PA.HFD.SAL$padj < 0.05, na.rm = TRUE)
write.csv(as.data.frame(SED.HFD.PGE_PA.HFD.SAL), file =
"/home/afb/AL3_data/AL3_SED.HFD.PGE_PA.HFD.SAL.csv")

#### 25)  SED.SD.SAL vs SED.HFD.PGE ######
SED.SD.SAL_SED.HFD.PGE <- results(ddsIF_3, contrast = c("group", "SEDSDSAL", "SEDHFDPGE" ))
SED.SD.SAL_SED.HFD.PGE <- SED.SD.SAL_SED.HFD.PGE[order(SED.SD.SAL_SED.HFD.PGE$padj),]
SED.SD.SAL_SED.HFD.PGE
summary(SED.SD.SAL_SED.HFD.PGE)
sum(SED.SD.SAL_SED.HFD.PGE$padj < 0.1, na.rm = TRUE)
sum(SED.SD.SAL_SED.HFD.PGE$padj < 0.05, na.rm = TRUE)
write.csv(as.data.frame(SED.SD.SAL_SED.HFD.PGE), file =
"/home/afb/AL3_data/AL3_SED.SD.SAL_SED.HFD.PGE.csv")

#### 26)  SED.SD.PGE vs SED.HFD.SAL ######
SED.SD.PGE_SED.HFD.SAL <- results(ddsIF_3, contrast = c("group", "SEDSDPGE", "SEDHFDSAL" ))
SED.SD.PGE_SED.HFD.SAL <- SED.SD.PGE_SED.HFD.SAL[order(SED.SD.PGE_SED.HFD.SAL$padj),]
SED.SD.PGE_SED.HFD.SAL
summary(SED.SD.PGE_SED.HFD.SAL)
sum(SED.SD.PGE_SED.HFD.SAL$padj < 0.1, na.rm = TRUE)
sum(SED.SD.PGE_SED.HFD.SAL$padj < 0.05, na.rm = TRUE)
write.csv(as.data.frame(SED.SD.PGE_SED.HFD.SAL), file =
"/home/afb/AL3_data/AL3_SED.SD.PGE_SED.HFD.SAL.csv")

#### 27)  PA.SD.SAL vs PA.HFD.PGE ######
PA.SD.SAL_PA.HFD.PGE <- results(ddsIF_3, contrast = c("group", "PASDSAL", "PAHFDPGE" ))
PA.SD.SAL_PA.HFD.PGE <- PA.SD.SAL_PA.HFD.PGE[order(PA.SD.SAL_PA.HFD.PGE$padj),]
```

```
PA.SD.SAL_PA.HFD.PGE
summary(PA.SD.SAL_PA.HFD.PGE)
sum(PA.SD.SAL_PA.HFD.PGE$padj < 0.1, na.rm = TRUE)
sum(PA.SD.SAL_PA.HFD.PGE$padj < 0.05, na.rm = TRUE)
write.csv(as.data.frame(PA.SD.SAL_PA.HFD.PGE), file =
"/home/afb/AL3_data/AL3_PA.SD.SAL_PA.HFD.PGE.csv")

#### 28)  PA.SD.PGE vs PA.HFD.SAL ######
PA.SD.PGE_PA.HFD.SAL <- results(ddsIF_3, contrast = c("group", "PASDPGE", "PAHFDSAL" ))
PA.SD.PGE_PA.HFD.SAL <- PA.SD.PGE_PA.HFD.SAL[order(PA.SD.PGE_PA.HFD.SAL$padj),]
PA.SD.PGE_PA.HFD.SAL
summary(PA.SD.PGE_PA.HFD.SAL)
sum(PA.SD.PGE_PA.HFD.SAL$padj < 0.1, na.rm = TRUE)
sum(PA.SD.PGE_PA.HFD.SAL$padj < 0.05, na.rm = TRUE)
write.csv(as.data.frame(PA.SD.PGE_PA.HFD.SAL), file =
"/home/afb/AL3_data/AL3_PA.SD.PGE_PA.HFD.SAL.csv")

#----------------------------------#------------------------------------------#------------------------------------------#
```
